# High-level correction of the sickle mutation amplified *in vivo* during erythroid differentiation

**DOI:** 10.1101/432716

**Authors:** Wendy Magis, Mark A. DeWitt, Stacia K. Wyman, Jonathan T. Vu, Seok-Jin Heo, Shirley J Shao, Fiona Hennig, Zulema G. Romero, Beatriz Campo-Fernandez, Matthew McNeill, Garrett R. Rettig, Yongming Sun, Patrick J. Lau, Yu Wang, Mark A. Behlke, Donald B. Kohn, Dario Boffelli, Mark C. Walters, Jacob E. Corn, David I.K. Martin

## Abstract

Sickle Cell Disease (SCD), one of the world’s most common genetic disorders, causes anemia and progressive multiorgan damage that typically shortens lifespan by decades; currently there is no broadly applicable curative therapy. Here we show that Cas9 RNP-mediated gene editing with an ssDNA oligonucleotide donor yields markerless correction of the sickle mutation in more than 30% of long-term engrafting human hematopoietic stem cells (HSCs), using a selection-free protocol that is directly applicable to a clinical setting. We further find that *in vivo* erythroid differentiation markedly enriches for corrected ß-globin alleles. Adoption of a high-fidelity Cas9 variant demonstrates that this approach can yield efficient editing with almost no off-target events. These findings indicate that the sickle mutation can be corrected in human HSCs at curative levels with a streamlined protocol that is ready to be translated into a therapy.

**ONE SENTENCE SUMMARY:** Cas9-mediated correction of the sickle mutation in human hematopoietic stem cells can be accomplished at curative levels.

## MAIN TEXT

SCD (*1*) is an attractive target for correction by gene editing(*2*–*5*) because hematopoietic stem cells (HSCs) can readily be harvested for *ex vivo* manipulation, and SCD is caused by a single point mutation in the ß-globin gene. Our strategy focused on xenografting of human stem and progenitor cells (HSPCs) to assess the edited genotype in the HSCs that drive long-term engraftment, assess the distribution of genotypes among engrafted cells, and examine globin expression in erythroid cells derived from edited HSCs (Figure 1B). Adult human peripheral blood CD34+ HSPCs homozygous for the sickle mutation were obtained by plerixafor mobilization. To correct the sickle mutation (Figure 1A), we electroporated HSPCs with a Cas9 ribonucleoprotein (RNP)(*6*) targeted to the site of the sickle mutation by a synthetic sgRNA with 3XMS protection (to improve stability of the guide RNA (*7*)), and a single stranded oligonucleotide donor template programming a wild type *HBB* allele. Our oligonucleotide donor facilitates homology-directed repair (HDR) that repairs the cleavage site using the donor sequence as template, alters the PAM sequence to prevent cleavage of the corrected allele, and changes the sickle mutation to the wild type sequence (Figure 1A). After electroporation, cells were injected into NOD.Cg-*Kit*^*W-41J*^ *Tyr*^+^ *Prkdc*^*scid*^ *Il2rg*^*tm1Wjl*^/ThomJ (NBSGW) mice, which permit engraftment of human HSCs and erythroid differentiation in the bone marrow (*8*, *9*). 16-20 weeks after injection, we analyzed human cells engrafted in the bone marrow (Figure 1B). We present data on 43 mice in 4 cohorts; mice in each cohort received aliquots of cells from a single electroporation. Human engraftment was robust, (52±27% mean±s.d., Figure 1C; Table S1). Immunophenotyping and colony assays indicated normal lineage potential of the xenografted cells (Figure S1).

**Figure 1.**
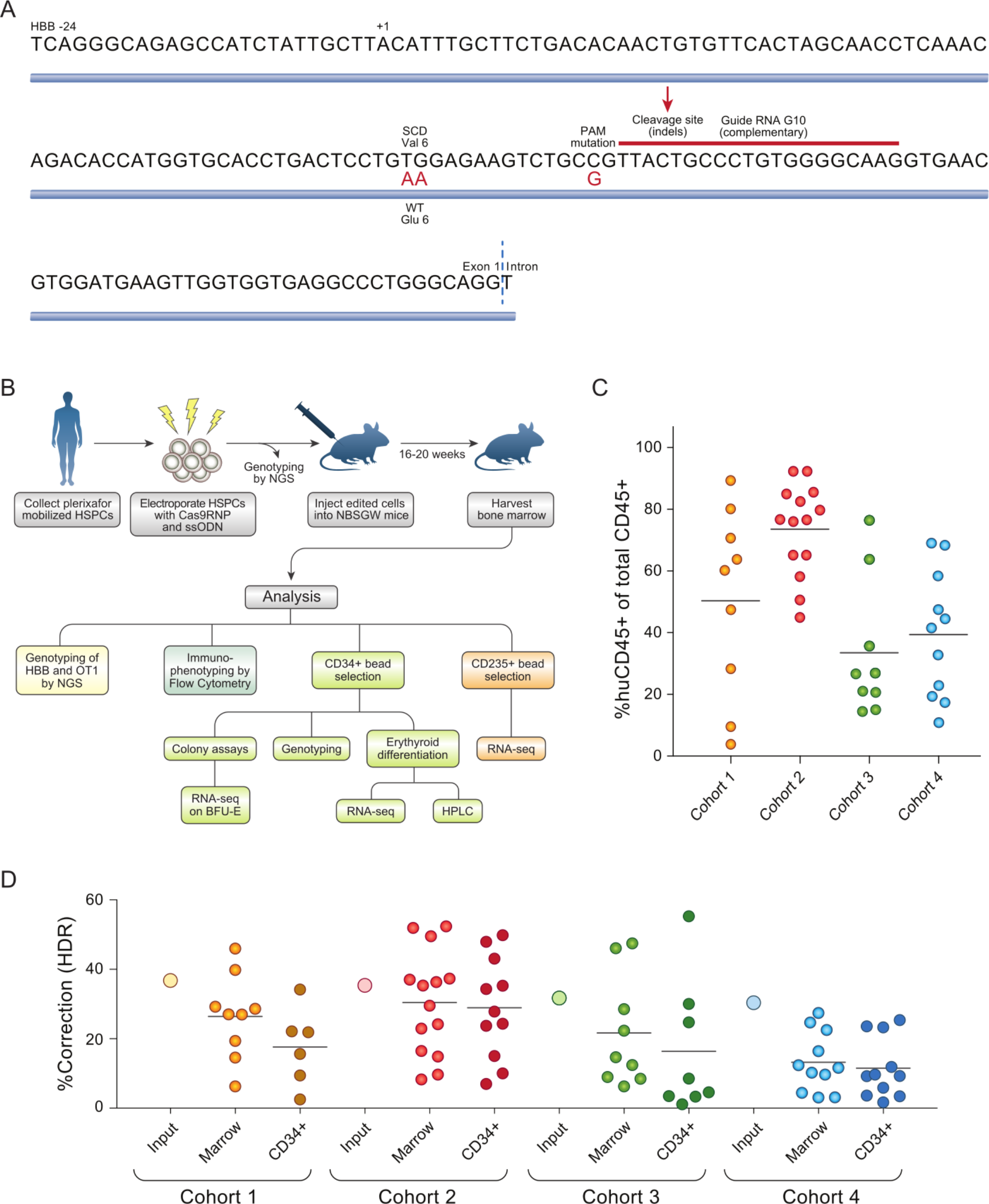
Correction of the sickle mutation in long-term xenografted hematopoietic cells. A) Schematic depicting targeting of the sickle mutation with the G10 guide RNA (in red) and the single-stranded DNA oligonucleotide donor CJ6A (blue). Sequence changes induced by the donor are shown in red; HDR proceeds from the Cas9 cleavage site (red arrow) but does not always extend to the site of the sickle mutation. B) Schematic outlining the large-scale xenografting experiment, and analysis of the engrafted cells. C) Engraftment of human hematopoietic cells in mouse bone marrow at 16-20 weeks post-injection, assessed by flow cytometry and displayed as percent of total marrow hematopoietic cells. D) Correction of the sickle mutation in xenografted marrow cells, and marrow CD34+ cells, at 16-20 weeks post-injection, expressed as percent of *HBB* alleles. The horizontal line denotes the average correction in each group; Input (large open circles) denotes percent correction in the pool of edited cells injected into each cohort.

We used PCR amplification of the edited region in *HBB* followed by next-generation sequencing (NGS)(*10*) to quantify editing in xenografted marrows (Figure 1D; Table S1), and found an average of 23.4% of alleles with the corrected genotype, and 65.2% with indels (Figure 1D; Figure S2; Table S1). Corrected alleles are distributed throughout the population of edited cells: the Hardy-Weinberg principle predicts that this allele frequency would result in >40% of cells carrying a corrected allele, but deviation from the Hardy-Weinberg distribution (see below and Figure S4) suggests a lower proportion that is still >30%; importantly, a single corrected allele is sufficient for normal erythrocyte function. Previous studies have found a dramatic decline in the proportion of HDR-corrected *HBB* alleles relative to input HSPCs (cells at the time of injection) (*2*, *4*, *5*), suggesting that HSCs carry out homology-directed DNA repair (HDR) less efficiently than the more committed progenitors that make up the majority of cells in the edited HSPC pool. However results in this experiment (Figure 1D) indicate that substantial levels of HDR can be achieved in HSCs.

Mice within the same cohort, despite receiving cells from the same electroporation, exhibit prominent variation in levels of gene correction (Figure 1D; Table S1). To investigate the basis of this variation, we examined the alleles present in xenografted marrow. Cas9 cleavage of *HBB* can lead to any of multiple alleles at the edited site: in the absence of HDR, repair by non-homologous end joining (NHEJ) introduces a variety of small insertions or deletions (indels). We determined the distribution of alleles at both the targeted *HBB* site and at a previously characterized intergenic off-target site (OT1) (*4*, *11*) (Figure 2; Figure S3).

**Figure 2.**
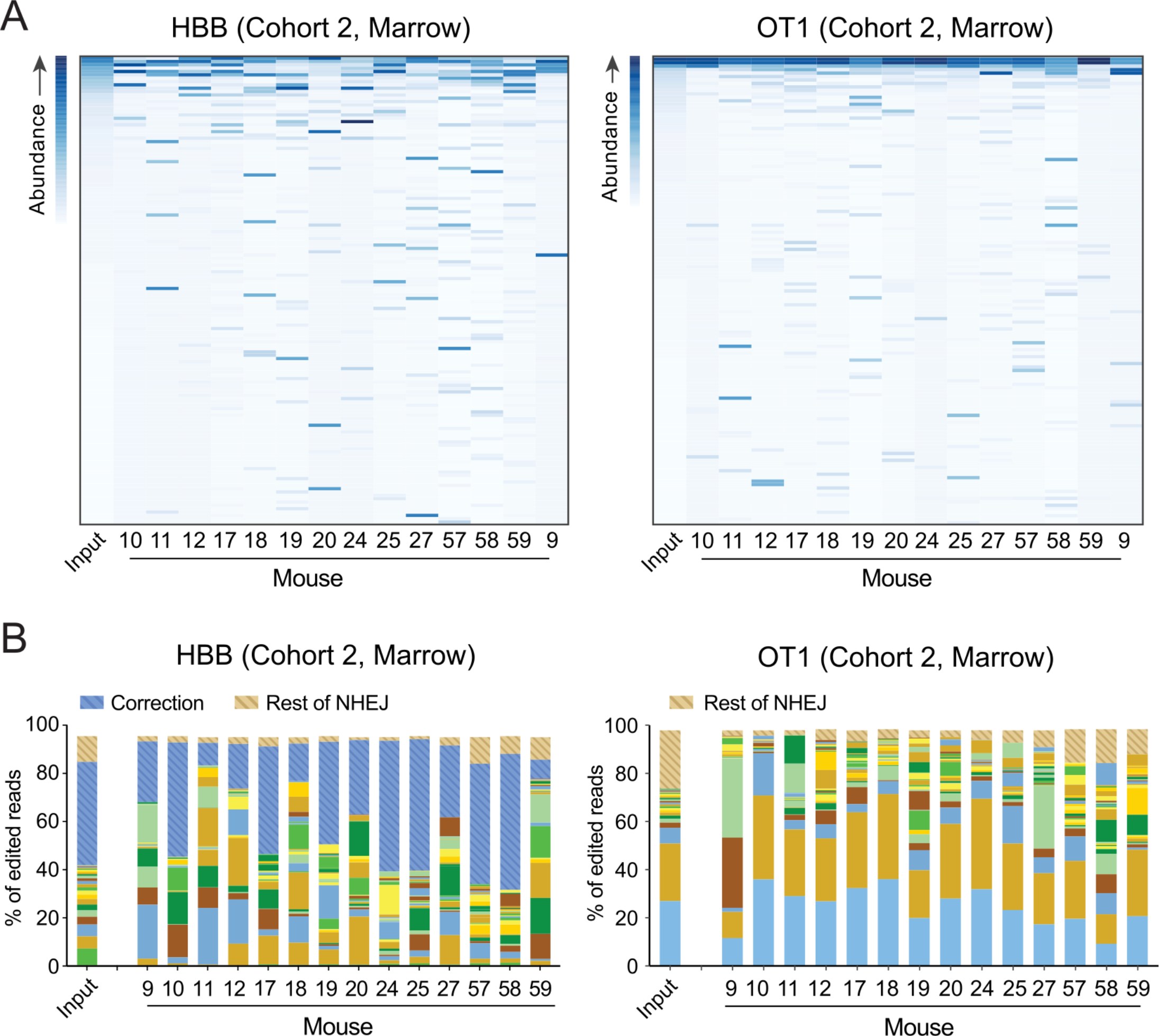
Analysis of allelic diversity in edited and xenografted hematopoietic cells. A) Heatmaps of abundance of alleles (rows) in input sample (first column) versus xenografted mice. Alleles are sorted (vertically) by decreasing abundance in the input sample, and filtered for alleles with at least 0.1% abundance in some mouse or the input, resulting in 140 HBB and 135 OT1 alleles. Note that the most common alleles in the input population are of minor abundance in some mice, while alleles that are rare in the input may be common in one or more mice. B) Stacked bar graphs indicating the contribution of the top 24 indel alleles (solid bars) at *HBB* (left) and OT1 (right) in each Cohort 2 mouse, as a fraction of all aligned and edited reads. All other NHEJ alleles are contained in brown striped bars (top segment); the corrected *HBB* alleles (HDR) are shown as a blue striped bar (second segment from top).

The edited HSPC pools prior to engraftment (“input”) contain a complex assortment of edited alleles. We found that each engrafted mouse carries a unique and smaller set of alleles, consistent with each receiving a discrete but still complex set of stem cells contributing to long-term engraftment. Alleles that are rare in the input population are prominent in some individual mice, but the identity of the prominent alleles differs among the mice (Figure 2; Figure S3). The outsize contribution of these alleles is consistent with some stem cells producing a much higher proportion of marrow cells than others. The observed mouse-to-mouse variation in sickle mutation correction may thus result from each mouse receiving a limited set of stem cells within which the corrected allele was represented to varying degrees, and by individual stem cell clones contributing disproportionately to the marrow cell population. Our results with gene-edited HSPCs are similar to observations of unique integration sites in lentivirus-transduced HSPCs (*12*, *13*).

We further characterized phenotypic and genotypic correction after engraftment in a subset of mice. Erythrocytes differentiated *in vitro* from immunoselected marrow CD34+ cells expressed corrected adult ß-globin mRNA and normal adult hemoglobin (HbA), elevated levels of γ-globin mRNA (HBG) and fetal hemoglobin (HbF), and markedly reduced levels of sickle ß-globin and sickle hemoglobin (HbS) (Figure 3A). To assess the distribution of *HBB* alleles within populations of xenografted cells, we inferred *HBB* genotypes using RNA-Seq data from 359 clonal BFU-E colonies derived from marrow CD34+ cells (Figure 3B; Figure S4). This reveals that a substantial proportion of HDR alleles are heterozygous with indel (NHEJ) alleles that are potentially equivalent to ß-thalassemia mutations. Wild type (corrected) *HBB* is dominant over sickle and null (indel) alleles: individuals heterozygous for the sickle mutation, or for a ß^0^ thalassemia, have functionally normal erythrocytes. The corrected (wild type) alleles in edited HSCs are distributed such that many erythrocytes have only a single wild type *HBB* allele. This distribution produces a proportion of functional erythrocytes that is higher than the proportion of corrected alleles observed in the population of CD34+ cells from which the erythroid colonies were derived (14.6% corrected alleles distributed among 24% functional erythrocytes). When this relationship is applied to the average gene correction over all xenografted mice (23.4%) it predicts a proportion of functional erythrocytes that is >30%.

**Figure 3.**
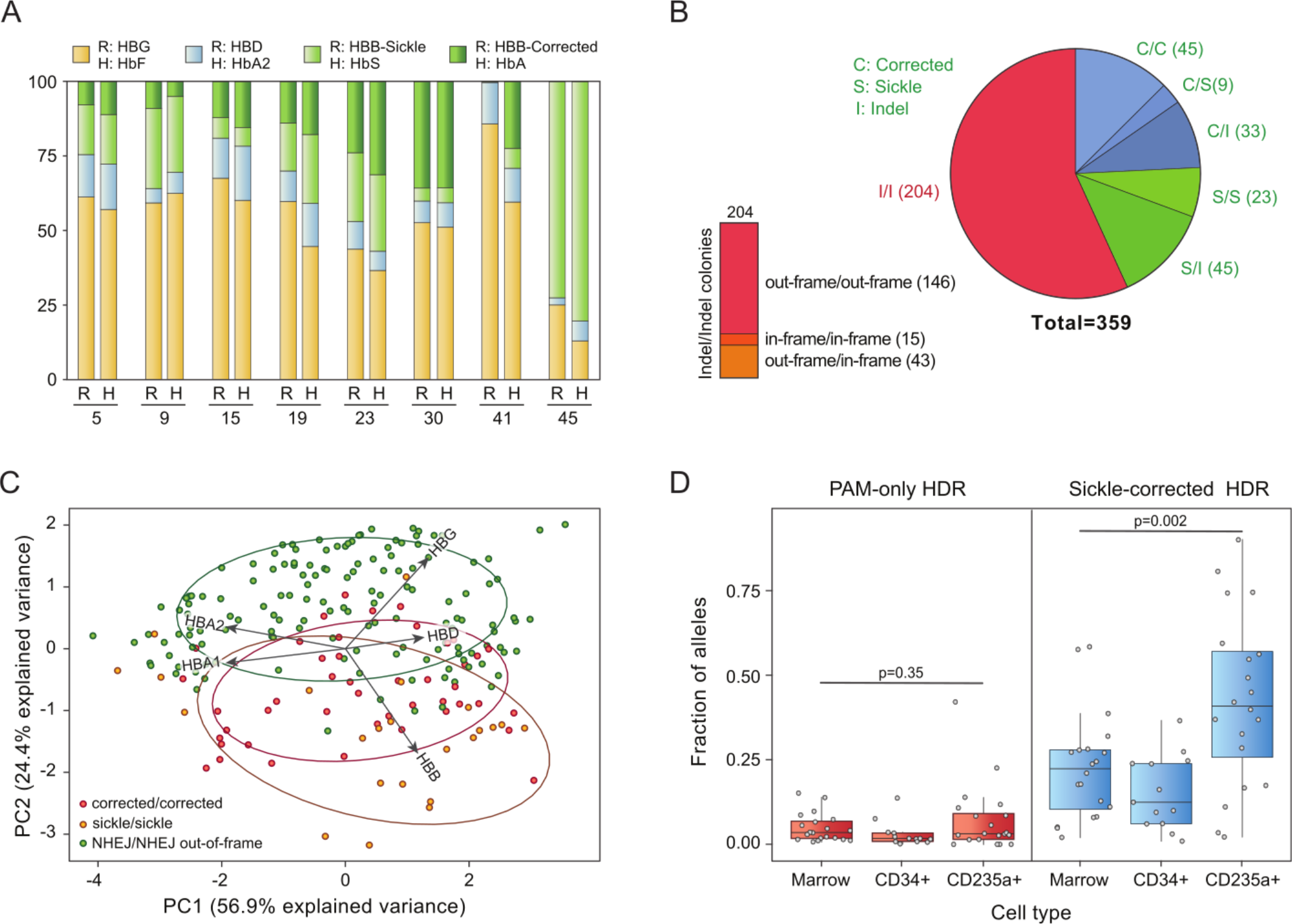
Globin expression, allelic assortment, and *in vivo* selection for corrected HBB genotype in erythroid cells. A) RNA-Seq (R) and HPLC (H) on erythrocytes differentiated from CD34+ marrow cells of 8 xenografted mice; mouse 45 was engrafted with unedited cells. For RNA-Seq, only ß-like globins are shown, in colors matching the corresponding hemoglobin assayed by HPLC. B) Genotypes inferred from RNA-Seq of clonal erythroid colonies differentiated from xenografted marrow CD34+ cells; homozygous NHEJ colonies are further characterized by in- vs. out-of-frame deletions (bar at left). C) Principal component analysis of RNA-seq data from erythroid colonies with homozygous genotypes: dots represent individual colonies; ellipses encompass 68% of colonies of each type. Colonies with out-of-frame indels at *HBB* have a reduced ratio of *HBB* to *HBG* mRNA. See also Figure S5. D) Distribution (box plot) of HDR allele frequency in total marrow, marrow CD34+ genomic DNA, and marrow erythroid (CD235+) cells; data for individual mice are shown as circles. A “sickle-corrected” HDR allele has both the PAM mutation and sickle correction, while a “PAM-only” HDR allele has the PAM mutation but no sickle correction. The corrected allele is enriched in erythroid cells compared to the PAM-only allele; see also Figure S6.

We assessed the effects of *HBB* genotype on the pattern of globin expression in the individual clonal erythroid colonies (Figure 3C). For this analysis, we excluded heterozygous colonies, and separated biallelic indels that alter the reading frame (which are likely to be equivalent to null ß-thalassemia mutations) from biallelic indels that retain the native reading frame. Principal component analysis on expression of the globin genes *HBA*, *HBG*, *HBD*, and *HBB* (Figure 3C, Figure S5) reveals that colonies with out-of-frame indels are skewed toward expression of *HBG*, the fetal ß-like globin gene, while those with in-frame deletions are closer to the pattern of colonies with either sickle or wild type (corrected) *HBB*. This analysis suggests that a large proportion of erythroblasts generated by edited HSCs have a ß-thalassemia phenotype, likely due to out-of-frame indels causing deficiency of *HBB* transcripts.

The findings discussed above (Figure 1, Figure 3B and 3C) indicate that our editing protocol produces a high proportion of indel alleles that are functionally equivalent to ß-thalassemia mutations. But in contrast to ß-thalassemia, the HSCs with non-functional *HBB* alleles are mixed with a high proportion of HSCs carrying corrected *HBB* alleles. Deficiency of ß-globin chains leads to apoptosis during late stage erythroid differentiation (“ineffective erythropoiesis”), and a similar phenomenon may occur in sickle cell disease(*14*, *15*).

Because cells carrying indels or the sickle mutation on both alleles may be subject to ineffective erythropoiesis, while cells with even one corrected *HBB* allele are functionally normal, we asked if erythroid cells in which either one or both *HBB* alleles had been corrected enjoy a selective advantage during erythropoiesis. NBSGW mice engrafted with human HSCs produce human erythroblasts in their bone marrow (*8*, *9*). We immunoselected human erythroid cells with antibody to CD235a (GlycophorinA), carried out RNA-Seq, and inferred the *HBB* genotypes present in the erythroblast population. For each individual mouse, we compared the proportion of corrected *HBB* alleles in the xenografted bone marrow, in CD34+ cells immunoselected from marrow, and in CD235a+ erythroblasts. To control for enrichment of alleles corrected by HDR, we distinguished two types of HDR: “PAM-only” HDR events mutate the Cas9 PAM without the conversion tract reaching the sickle SNP; whereas “sickle-corrected” HDR events mutate the Cas9 PAM and also correct the sickle SNP (Figure 1A). We observe a marked enrichment of “sickle-corrected” HDR alleles in erythroblasts when compared to marrow or CD34+ cells from the same mouse, but no enrichment of “PAM-only” HDR alleles (Figure 3D, Figure S6). This observed enrichment of corrected *HBB* alleles in edited erythroblasts may reflect the loss of sickle, ß-thalassemia, and sickle/ß-thalassemia cells *in vivo*.

While Cas9 RNP/ssDNA-based gene correction may provide an effective therapeutic option for SCD, wild type Cas9 complexed with the G10 sgRNA exhibits off-target cleavage activity at the previously-characterized site OT1 (*4*, *11*), where after engraftment an average of 47.5% of alleles have indels (Figure S2). While indels at OT1 have no known or predicted deleterious effect, cleavage at other genomic sites has the potential to promote neoplastic progression. To identify and characterize additional potential off-targets, we developed an exhaustive strategy that combines GUIDE-seq (*16*), unbiased off-target identification, bioinformatic searches, and pooled-primer quantification. GUIDE-seq in K562 cells revealed *HBB* and OT1, and an additional five off-target sites; none of the five sites are in a coding region (Figure 4A, Table S2). We term the most prevalent of these newly identified sites “OT2”. GUIDE-seq in HSPCs found only the OT1 site (Figure S7).

**Figure 4.**
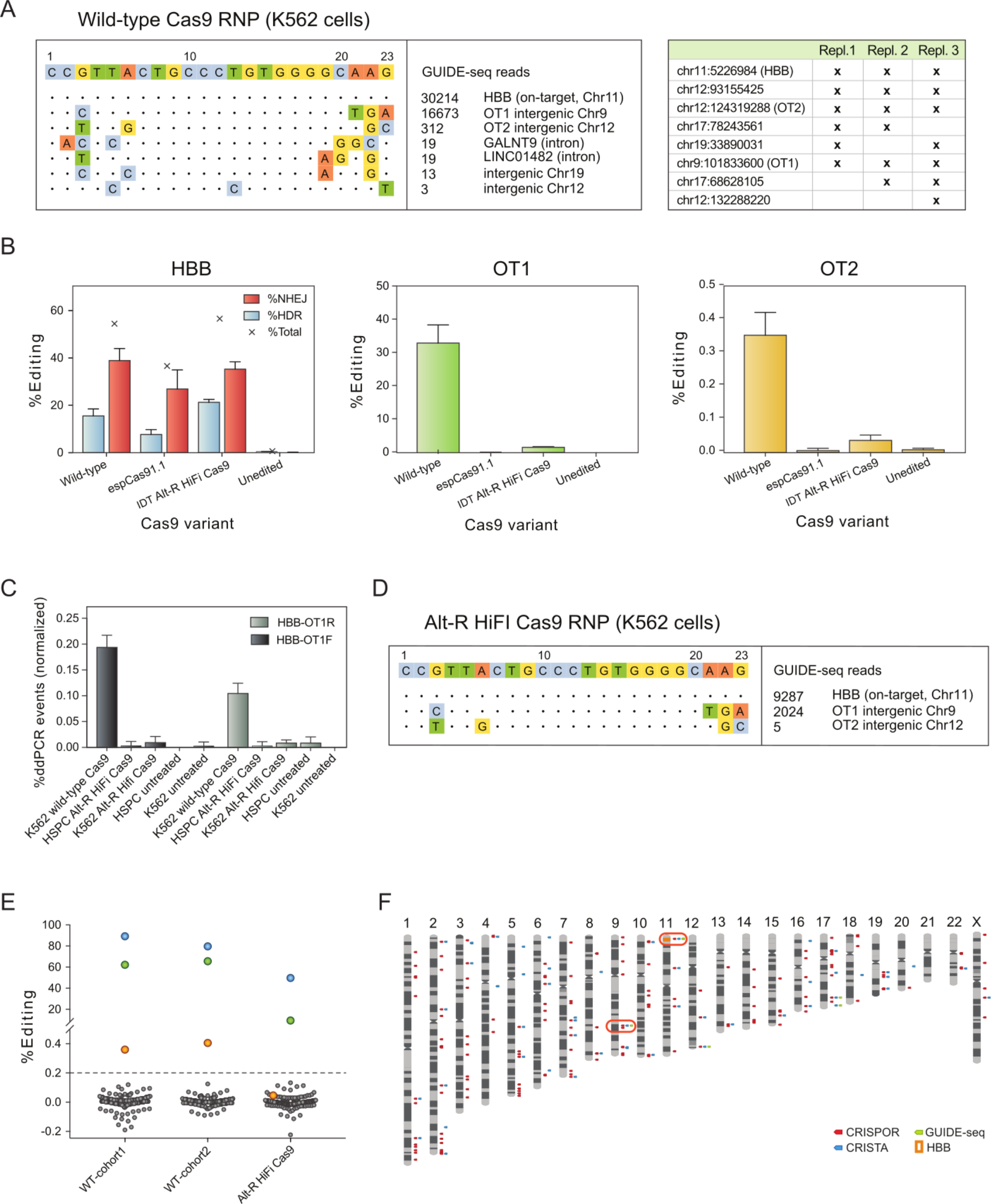
Assessment of off-target cleavage. A) Representative GUIDE-seq with the Cas9 RNP in K562 cells (left), and GUIDE-seq hits detected in three independent replicates (table at right). B) Comparison of editing at *HBB* by high-fidelity Cas9 variants espCas9-1.1 (*19*) and Alt-R HiFi Cas9 (*18*) in healthy donor HSPCs. C) Droplet digital PCR (ddPCR) detects translocation linking the region upstream of *HBB* with regions either upstream (OT1F) or downstream (OT1R) of OT1, when *HBB* is targeted with wild-type Cas9, but not with Alt-R HiFi Cas9. D) Representative GUIDE-seq with Alt-R HiFi Cas9 RNP in K562 cells. E) Total gene editing rates (%HDR + %NHEJ) measured by pooled-primer PCR at 190 of 201 identified off-targets, in the edited HSPCs injected into Cohorts 1 and 2 (“input” in Figure 1C), and in healthy donor HSPCs edited with Alt-R HiFi Cas9; indels observed at the same sites in untreated cells are subtracted. Blue dots: on-target *HBB*; green dots: OT1; orange dots: OT2. Editing at >0.2% of alleles (dashed line) by wild-type Cas9 is observed only at OT1 and OT2, and by high-fidelity Cas9 only at OT1. F) Off-target sites identified by GUIDE-seq, CRISTA, and CRISPor; sites where editing was detected (in >0.2% of alleles using high-fidelity Cas9) are indicated with a red box.

Off-target cleavage may be mitigated by use of recently developed high-fidelity Cas9 variants. We compared targeting of *HBB* and OT1 by RNPs made up of the 3XMS-G10 guide with either WT Cas9, espCas9-1.1(*17*), HF-1(*18*), or Alt-R HiFi Cas9(*19*). In K562 cells (Figure S8), all three high fidelity variants reduced off-target editing at both OT1 and OT2. However, HF-1 reduced on-target editing at *HBB* and was therefore not tested further. In HSPCs (Figure 4B), only the AltR HiFi variant (IDT) maintained high levels of HDR editing at *HBB* while reducing indel formation at OT1 by ~20-fold, and OT2 by ~10-fold (Figure 4B, Figure S8). We therefore selected the AltR HiFi variant for further study. While indels at OT1 are unlikely to be deleterious, translocations are more of a concern. Translocations between *HBB* and OT1 are detectable in K562 cells and CD34+HSPC edited with wild-type Cas9, but undetectable with AltR HiFi Cas9 using a droplet digital PCR assay(*20*) (Figure 4C). GUIDE-seq with AltR HiFi Cas9 RNP in K562 cells revealed only *HBB*, OT1, and OT2 (Figure 4D), and GUIDE-seq events were reduced for OT1 and OT2 compared to WT Cas9.

For bioinformatic identification of candidate off-target sites we used CRISPor (*21*), which relies on sequence similarity, and CRISTA (*22*), which relies on machine learning. These tools produced a list of 201 potential off-target sites (Figure 4F). We quantified editing in HSPCs at all 201 sites using a pooled-primer PCR approach capable of simultaneous amplification (*19*). Wild type Cas9 edited only OT1 and *HBB* in >1% of alleles (Figure 4E, Table S3), but some off-target activity was seen at OT2 (~0.2% of alleles). Alt-R HiFi Cas9 reduced editing at OT1 to 2.1% of alleles, and editing at OT2 was reduced below the limit of detection (from 0.4% to 0.03%, Figure 4E, Table S3). No editing was observed at the 199 other sites with HiFi Cas9. We conclude that, of the 201 potential off-target sites (Figure 4F), only two (OT1 and OT2) are *bone fide* off targets of the wild-type 3xMS-G10 Cas9 RNP, and only one (OT1) is an off-target of the AltR HiFi 3xMS-G10 Cas9 RNP.

Two problems have been presented by attempts to correct the sickle mutation in HSCs: low levels of correction in long-term engrafted cells, which might reflect a low level of HDR competence in HSCs, and the creation of *HBB*-null (ß-thalassemia) alleles by NHEJ repair of Cas9 cleavage. We observe correction of the sickle mutation in more than 20% of ß-globin alleles, distributed among more than 30% of long-term engrafting HSCs, demonstrating that HSCs are competent to carry out HDR at levels sufficient to cure SCD, ß-thalassemia, and many other genetic blood disorders. Evidence from mixed donor chimerism after allogeneic hematopoietic stem cell transplantation for SCD and ß-thalassemia indicates that that as little as 10% (SCD) or 20% (ß-thalassemia) of donor cells are sufficient to render a recipient free of disease, because functional erythrocytes (carrying at least one wild type *HBB* allele) have a selective advantage during erythropoiesis and in circulation (*23*–*26*). We present evidence that the corrected *HBB* allele enjoys a selective advantage *in vivo* over *HBB*-null (ß-thalassemia) alleles created by NHEJ in our editing protocol, thus amplifying the effect of gene correction. Our streamlined protocol does not involve any selection, and produces robust multiclonal engraftment with little off-target editing. It is thus suitable for translation into a therapy based on correction of the sickle mutation in autologous HSCs, which has the potential to cure SCD while avoiding complications inherent in other therapies.

## ACKNOWLEDGMENTS

We thank Larry Young and Flora Ting for performing HSPC injections, Sean Xu for analysis of single-colony RNA-Seq, and Alicia Garcia and Marci Moriarty for their assistance in procuring sickle HSPCs.

## Data and materials availability

All sequence data has been deposited in the NCBI Sequence Read Archive under BioProject number PRJNA498110.

## SUPPLEMENTARY MATERIALS

Materials and Methods

Figures S1 – S8

Tables S1 – S3

References (27-31)

