## Supplementary Material for "High-level correction of the sickle mutation amplified *in vivo* during erythroid differentiation"

### SUPPLEMENTARY INFORMATION

#### Materials and Methods

*Gene editing reagents.* sgRNA G10 was characterized in our previous work(*4*). In this study all sgRNA was synthetic, carried the 3X-MSP modification(*7*), and was obtained from Synthego, Trilink, or Agilent. Wild-type Cas9 protein, Cas9 HF-1, and Cas9 espCas9-1.1 were purified according to published protocols (*6*). AltR HiFi Cas9 was purchased from IDT, Inc. or purification tag-free from Aldevron, Inc. We found no difference in editing outcomes between IDT and Aldevron high fidelity Cas9. ssDNA CJ6A (IDT) was synthesized using the Ultramer synthesis platform.

*CD34+ Cells.* CD34+ HSPC homozygous for the sickle mutation from clinical trial participants mobilized with plerixafor were obtained with consent from apheresis discard material at Children's Hospital Oakland, purified by Allcells, Inc., and cryopreserved in aliquots. Healthy donor CD34+ cells were purchased from AllCells.

*Gene editing protocol for CD34+ HSPC.* Cryopreserved CD34+ cells were thawed according to AllCells instructions, and cultured for 2 days in StemSpan SFEM with CC110 supplement (StemCell Technologies). For each cohort of mice, cells were thawed, combined, and cultured together, electroporated in $10^6$ cell aliquots with RNP/ssDNA from a master mix, then recombined and cultured together before injection. Just prior to electroporation, cells were pelleted at 100 x g for 10 minutes, and resuspended to 1-3x$10^4$ cells/µL in Lonza P3 buffer; during the electroporation procedure, cells did not remain in P3 for longer than 20 minutes. While cells were in the centrifuge, the Cas9 RNP/ssDNA mixture was prepared (10.6 µM sgRNA (3xMS-G10 sgRNA), 8.8 µM Cas9 protein, and 11.8 µM ssDNA CJ6A (or CJ6 when editing healthy donor HSPC) in Cas9 RNP buffer (20 mM HEPES pH 7.50, 150 mM KCl, 1 mM $MgCl_2$, 10% glycerol, 1 mM TCEP) as described (*6*)). The RNP/ssDNA was mixed with cells at a 0.375:1 ratio (i.e. 30 µL RNP/ssDNA mix to 80 µL CD34+ cells). The mixture was placed in a Lonza Nucleofector cuvette (20 µL "S" or 100 µL "L") and electroporated using Lonza electroporation code ER100 on a Lonza 4D Nucleofector. Immediately after electroporation, at least 2 volumes of SFEM/CC110 was layered on top of cells for 5-10 minutes before gently transferring to a culture dish. Cells were cultured in SFEM/CC110, overnight before injection into mice (below), or 5-7 days before genotyping by next-generation sequencing. Prior to injection, cells were pelleted and resuspended in PBS.

*Xenografting of human CD34+ HSPCs into NBSGW mice*. NBSGW mice (JAX 026622) were maintained in clean conditions. NBSGW mice have the immunodeficient phenotype of NSG mice, but will accept human hematopoietic stem cell grafts without prior irradiation(*8*). 7-8-week-old female (cohorts 1 and 2) or male (cohorts 3 and 4) mice were injected with 6-7x$10^5$ edited cells in 200µL PBS via the lateral tail vein. Mice were observed, but subjected to no further procedures until euthanasia at 16-20 weeks after injection. All injected mice survived until that point without evidence of morbidity. After euthanasia, bone marrow (femur) was recovered for analysis.

*Genotyping of edited cells by next-generation sequencing.* Genomic DNA from cells in tissue culture were extracted in QuickExtract solution (Epicentre, Inc.), to a density of >2,000 cells/µL, according to manufacturer's instruction. Genomic DNA from xenografted bone marrow or bone marrow-derived CD34+ cells was extracted using the Machery-Nagel Nucleospin Blood kit in a 96 well format, according to manufacturer's instruction. PCR amplicons from either *HBB* or OT1 were generated as previously described(*4*), except that in the second PCR a short stub

(GCTCTTCCGATCT) was added to the 5' end of both primers, to match the sequence of custom-designed amplify-on Illumina sequencing adaptors, which were ligated to the second PCR amplicon through a third short-cycle (15 cycle) PCR using GXL polymerase and manufacturer's recommended cycling temperatures. The resulting amplicons were pooled and sequenced on an Illumina MiSeq using the 600 cycle v3 kit and a 2x300 paired-end sequencing read.

*Flow cytometric analysis of xenografted cells*. Cells were flushed from bone marrow with PBS, then prepared by passage through a 21G needle, filtering in a 40μm cell strainer to create a single-cell suspension, and red cell lysis with Qiagen EL buffer. Cells were stained with antibodies to the indicated cell surface markers, and analyzed on a BD FACS Fortessa flow cytometer. Flow cytometry data were analyzed with FlowJo. Antibodies, all from BD Pharmingen, were: APC Rat anti-Mouse CD45(561018, clone 30-F11), FITC Rat anti-Mouse CD45(553079, clone 30-F11), FITC Mouse anti-Human CD45 (555482, clone HI30), V450 Mouse anti-Human CD45 (560367, clone HI30), BV421 Mouse anti-Human CD3 (563798 clone SK7), BV421 Mouse anti-Human CD56 (562752, clone NCAM16.2), V450 Mouse anti-Human CD19 (644491, clone SJ25C1), FITC Mouse anti-Human CD33 (561818, clone HIM3-4), BV421 Mouse anti-Human CD34 (562577, clone 581).

*Immunoselection of CD34+ or Glycophorin A+ cells from xenografted marrows*. Bone marrow cells intended for immunoselection were not subjected to red cell lysis. Cells were immunoselected with MACS microbeads according to the manufacturer's instructions (CD34: Miltenyi 130-046-702; Glycophorin A: Miltenyi 130-050-501). CD34+ cells were further enriched by loading the eluate from the first column onto a second column and repeating the separation procedure.

*CFU progenitor assay in methylcellulose* Following immunoselection of CD34+ cells, cells were cultured for 3 days in StemSpan SFEM with CD34 expansion supplement (StemCell Technologies). Cells were then plated in Methocult Express (StemCell Technologies) at a density of 4-500 cells per 35mm well in a 6-well SmartDish (StemCell Technologies). Following ~14 days in culture, colonies were identified and counted under a microscope.

*Differentiation of CD34+ cells into erythroblasts*. Following immunoselection, CD34+ cells were transferred to StemSpan SFEM medium with StemSpan CD34 Expansion supplement (StemCell Technologies) and cultured for 3 days. For erythroid colonies, cells were then plated onto Methocult Express (StemCell Technologies); after ~14 days, erythroid colonies were identified by microscopy and picked individually for RNA-Seq analysis. For bulk erythroid culture (HPLC and RNA-Seq) expanded CD34+ cells were transferred to SFEM II medium with Erythroid Expansion supplement (StemCell Technologies) and grown for 8-10 days with maintenance of optimal density (200,000-1,000,000 cells/mL). The resulting erythroid progenitors were transferred to SFEM II with 4U/ml erythropoietin (Life Technologies), 3% normal human AB serum (Sigma), and 1 μM mifepristone (Sigma). They were cultured for a further 5-6 days with daily monitoring of cell density and morphology, with cell density maintained below $10^6$ cells/mL. For HPLC, cells were then lysed in hemolysate reagent (Helena Laboratories) for preparation of hemoglobin; for RNA-Seq, cells were extracted with the Direct-zol RNA Kit (Zymo Research).

*RNA-seq analysis of edited SCD HSCs.* Total RNA from marrow-isolated cells or SCD erythroblasts (~$5x10^6$, differentiated *in vitro* as described above) was isolated with the Direct-zol RNA Kit (Zymo Research). RNA integrity was checked on a Fragment Analyzer (Advanced Analytical); cDNA was synthesized from this RNA following the Smart-seq2 method (*27*), and fragmented with the Covaris apparatus. From the Covaris fragments, indexed sequencing libraries were constructed with the ThruPlex DNA-seq kit (Rubicon Genomics) and sequenced on an Illumina HiSeq 4000 sequencer for 50 cycles (single read) at the Berkeley GSL. Resulting RNA-seq reads for each sample were quantified using the program kallisto 0.43.1 (*28*), against a reference sequence consisting of the main isoforms of each globin gene at the alpha- and beta-globin loci. The following Ensembl transcript IDs were used: ENST00000320868.9 (HBA1), ENST00000251595.10 (HBA2), ENST00000199708.2 (HBQ1), ENST00000356815.3 (HBM), ENST00000354915.3 (HBZP1), ENST00000252951.2 (HBZ), ENST00000335295.4 (HBB), ENST00000380299.3 (HBD), ENST00000454892.1 (HBBP1), ENST00000330597.3 (HBG1), ENST00000336906.4 (HBG2), ENST00000292896.2 (HBE1). Kallisto returns the relative abundance of each mRNA in transcripts-per-million. Expression of each gene is reported as the proportion of expression of all beta-like genes (HBB, HBD, HBG1, HBG2).

Globin transcripts have high levels of sequence identity; to evaluate the ability of kallisto to correctly assign a read to the transcript from which it was derived, we carried out simulations using computationally generated 50-base reads. The globin transcripts listed above were mixed in known proportions that simulate relative transcript levels in erythrocytes (HBA-1: 0.7, HBA-2: 0.7, HBM: 0.01, HBB: 1.00, HBD: 0.04). HBG-1 and HBG-2 were varied between 0.005 and 0.10 to simulate the effect of varying levels of gamma-globin expression in a mixture. Sequence segments of 50 nucleotides in length were sampled randomly from this mixture and from its reverse complement (number of samples = 1,000,000), and aligned as described above. The random sampling procedure was repeated 10 times for each mixture. For each alignment result, we calculated the mean ratio of observed over expected counts for each transcript, which we used as adjustment factors for the counts reported by kallisto after aligning real data; the difference in transcript counts before and after adjustment ranged from -1% to 0.5%. The computed adjustment factors are listed in the table below:

| | |
|---|---|
| HBA-1 | 1.0398 |
| HBA-2 | 0.8891 |
| HBM | 0.8195 |
| HBB | 1.0402 |
| HBD | 1.1146 |
| HBG-1 | 1.3218 |
| HBG-2 | 1.0079 |

Analysis of haplotype and genotype frequencies from RNA-seq data was performed by first aligning the reads to the HBB transcript with BowTie2 (*29*) followed by analysis of the alignments with FreeBayes (*30*). Populations of cells isolated from the marrow (marrow, $CD34^+$, and $CD235a^+$) were analyzed with the option `pooled-continuous` to obtain haplotype frequencies; single colonies of SCD erythroblasts were analyzed to obtain genotypes of the region containing the sickle site and the PAM motif.

*HPLC analysis of hemoglobin in erythroid cells derived from xenografted marrow*. HPLC was carried out on extracts from bulk-differentiated erythroblasts (above) and data was analyzed as previously described (*9*, *10*).

*Analysis of genome editing events from next-generation sequencing data*. For each mouse sample and input sample per mouse cohort, targeted amplicon deep sequencing was performed both at the cut site and at OT1 and OT2, the primary off-target sites, as indicated. Paired end reads for each target (typically sequenced to a depth of at least 15,000 reads per sample, with an average of

~114,000 reads per sample) were analyzed based on a reimplemented version of the CRISPResso program(*10*). Briefly, paired end reads are trimmed to remove adapters and poor-quality trailing bases and then paired end reads are joined into a single read. Given the HBB reference sequence and the donor ssODN sequence plus the guide, each of the three HDR locations in the donor sequence is assessed for every read and then percent editing is output for each position. NHEJ percent is tallied for any read with an indel within a 6 basepair window around the cut site. For OT1, when no donor sequence is given, just NHEJ is assessed for the target.

*Analysis of allele identity and representation before and after engraftment in NBSGW mice (Figure 2).* Once all the reads have been classified for both targets in marrow and CD34+ samples for all mice (where available), an allelic abundance matrix is constructed for each cohort and target location for bone marrow and CD34+ cells. The columns are samples (mice in the cohort or input sample for the cohort) and rows in the matrix are alleles, represented by a 40 basepair window around the cut site and the values are the percent of aligned reads that that allele represents in the mouse or input sample. The resulting matrix contains the abundance for all alleles in every sample. This matrix is used as input for constructing a heatmap for each cohort for each target site in marrow and CD34+ cells where rows are alleles, columns are mice or the input sample, and the intensity represents the abundance of the allele. The alleles are sorted from top to bottom by abundance in the input sample, and normalized by column. Alleles which do not have greater than 0.1% abundance in any column are filtered out. Figure 2C-D is constructed by extracting out the top 24 alleles in each mouse in a cohort and the input sample and the abundance of those shared alleles across mice.

*Identification of off-targets using CRISTA, CRISPor, and GUIDE-seq.* The list of off-target loci in the genome was constructed using several overlapping methods. One *in vivo* method, two *in silico* methods and a direct homology search were employed and then the results were combined. GUIDE-Seq(*16*) is an *in-vivo*assay that identifies off target loci by inserting a dsODN sequence into double-stranded breaks created by the guide. The two *in silico* prediction methods used were CRISPOR(*21*) that primarily relies on sequence similarity to the guide sequence (allowing up to six mismatches between the guide and the predicted target) and CRISTA(*31*) that uses a machine learning approach incorporating many diverse features into its model. Finally, a BLAST homology search was performed with the ssODN sequence and the AAV sequence against the human genome sequence to identify possible integration sites. Combining the loci from all the methods resulted in a master list of 201 putative off-target sites.

*GUIDE-seq in K562 cells*. GUIDE-seq was adapted for use with Cas9 RNP from the published protocol(*16*). Briefly,100,000-200,000 K562 cells were cultured in IMDM supplemented with 10% fetal bovine serum, sodium pyruvate, and penicillin/streptomycin to mid-log phase ($0.5\text{-}1.0 \times 10^6$ cells/mL) before electroporation with 90 pmol Cas9 RNP assembled as described (*10*), with the addition of 100 pmol GUIDE-seq dsDNA oligonucleotide, using a Lonza 4d electroporator and a 20 µL cuvette. 48 hours after electroporation, cells were harvested and prepared for GUIDE-seq using the published protocol, with the addition of 5 PCR cycles in both PCR1 and PCR2. GUIDE-seq analysis and visualization used the published code (https://github.com/aryeelab/guideseq).

*Pooled primer PCR assay design (IDT).* To examine Cas9 activity at each target, we used custom software to design a pool of multiplex rhPCR, blocked-cleavable primers(*19*). Primer pairs were designed for each target; targets were 17-31nt. Primers were placed at least 20nt away from each target and formed amplicon inserts <215nt in unedited GRCh38 DNA. Optimal pairs were selected for the multiplex PCR reaction such that they would be unlikely to form primer dimers. Multiplex PCR primers, PCR mastermix, and associated reagents were provided by Integrated DNA Technologies.

*Pooled primer PCR.* The assay requires at least 10 ng of genomic DNA (gDNA). gDNA from edited HSPC was extracted using the Qiagen Blood and Tissue Kit. For each sample, two PCRs were performed, one with primers for 195 targets, and one with primers for 5 targets. The first PCR consisted of 10 cycles, followed by 1.5X SPRI bead clean-up using the pooled primer mix to amplify. The second (indexing) PCR consisted of 23 cycles using IDT rhAmpSeq indexing primers P5 and P7, followed by SPRI clean-up. The resulting PCRs were pooled and sequenced using an Illumina HiSeq, 2x150 paired end read. Gene editing events were assessed at each site by alignment of the data to human genome assembly hg38. The results, including off-target coordinates, are presented in Table S3.

*Droplet Digital PCR for translocations.* Droplet digital PCR for translocations between *HBB* and OT1 was performed as in Long *et al*(*20*) using the same forward and reverse primers and probe for *HBB,* a primer for OT1 upstream of the target site (OT1F, ctgaggaggaaacacataatgagagt), a primer for OT1 downstream of the target site (OT1R, gcggtggctctcaaatatcaatc), along with the published probe and primers for the untargeted positive control site in UC378. Each genomic DNA sample was used as template for 8 reactions for each of three primer pairs: HBB(f)-HBB(r), HBB(f)-OT1(f) and HBB(f)-OT1(r). For each sample the ratio of events for the indicated amplicon to UC378 control amplicon was calculated. For each sample, the translocation frequency was taken as the ratio for the HBB-OT1(f or r) samples divided by the ratio for the HBB(f)-HBB(r) samples.

### FIGURES

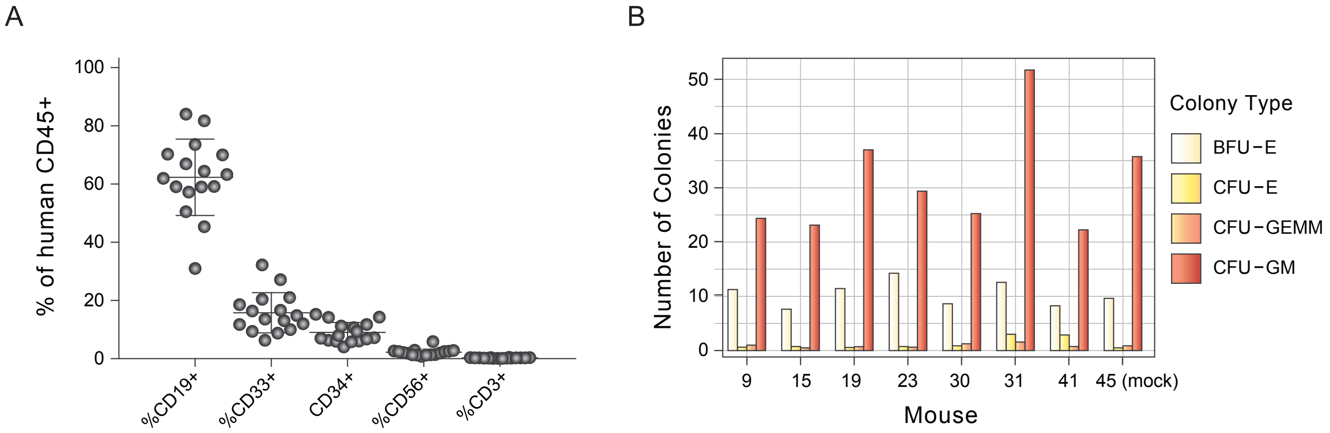

**Figure S1.** Multilineage differentiation of xenografted human hematopoietic cells. A) Immunophenotyping of human cells in mouse marrow by flow cytometry for CD19 (B cells), CD33 (myeloid cells), CD34 (progenitors), CD56 (NK cells), and CD3 (T cells). B) Colony assays seeded by human CD34+ cells isolated from mouse marrows after 16-20 weeks engraftment.

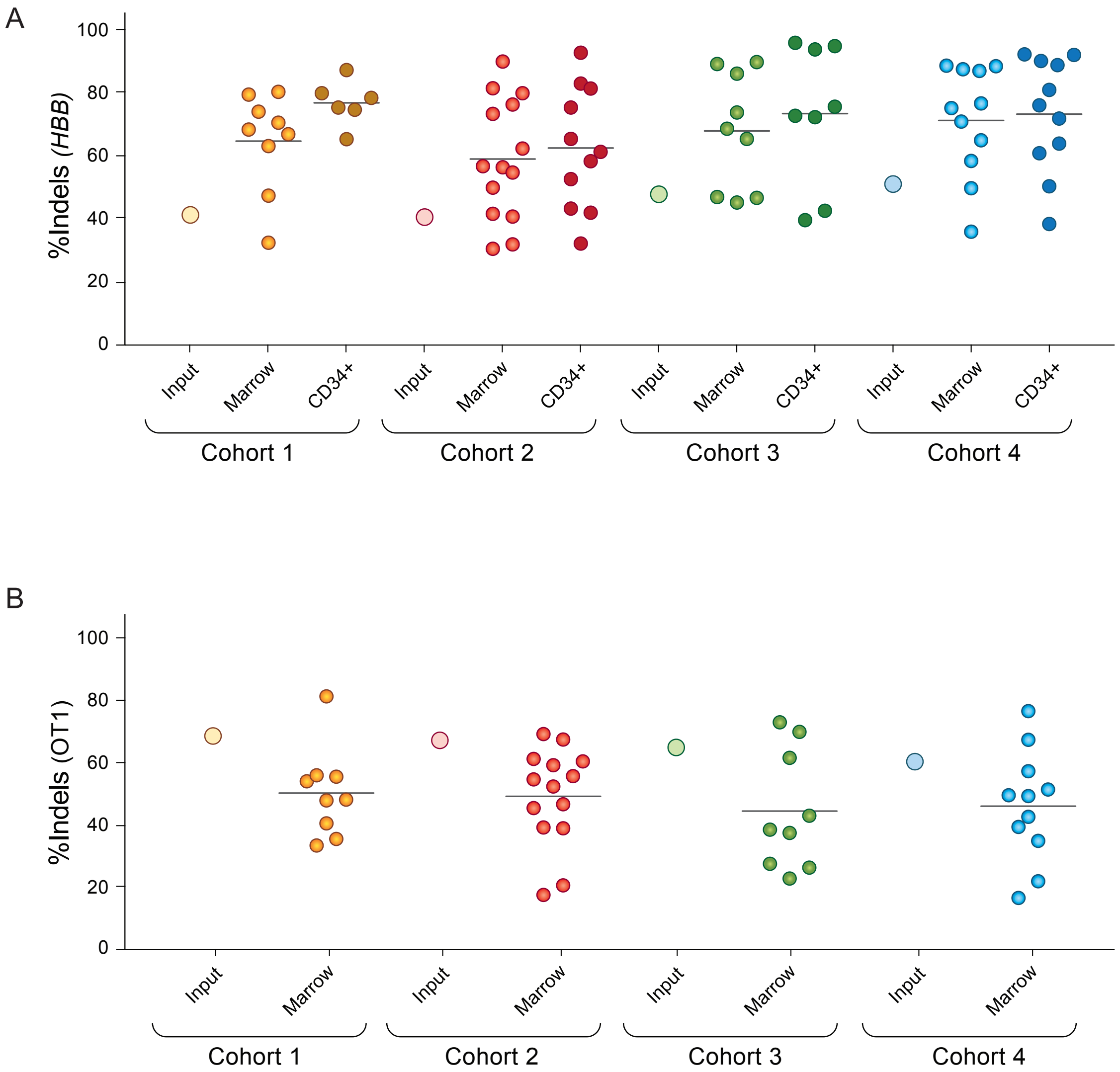

**Figure S2. NHEJ creates indels at the HBB target site and the off-target OT1 site. A.** Indels at the *HBB* target site in xenografted marrow cells, and marrow CD34+ cells, at 16-20 weeks post-injection, expressed as percent of *HBB* alleles; related to Figure 1D and Table S1. The horizontal line denotes the average indel percent in each group; Input (large open circles) denotes the percent in the pool of edited cells injected into each cohort. B. Indels at intergenic off-target OT1 (chr9:101833584-101833606 in human genome assembly hg38) in xenografted marrow cells (labeled and displayed as for A).

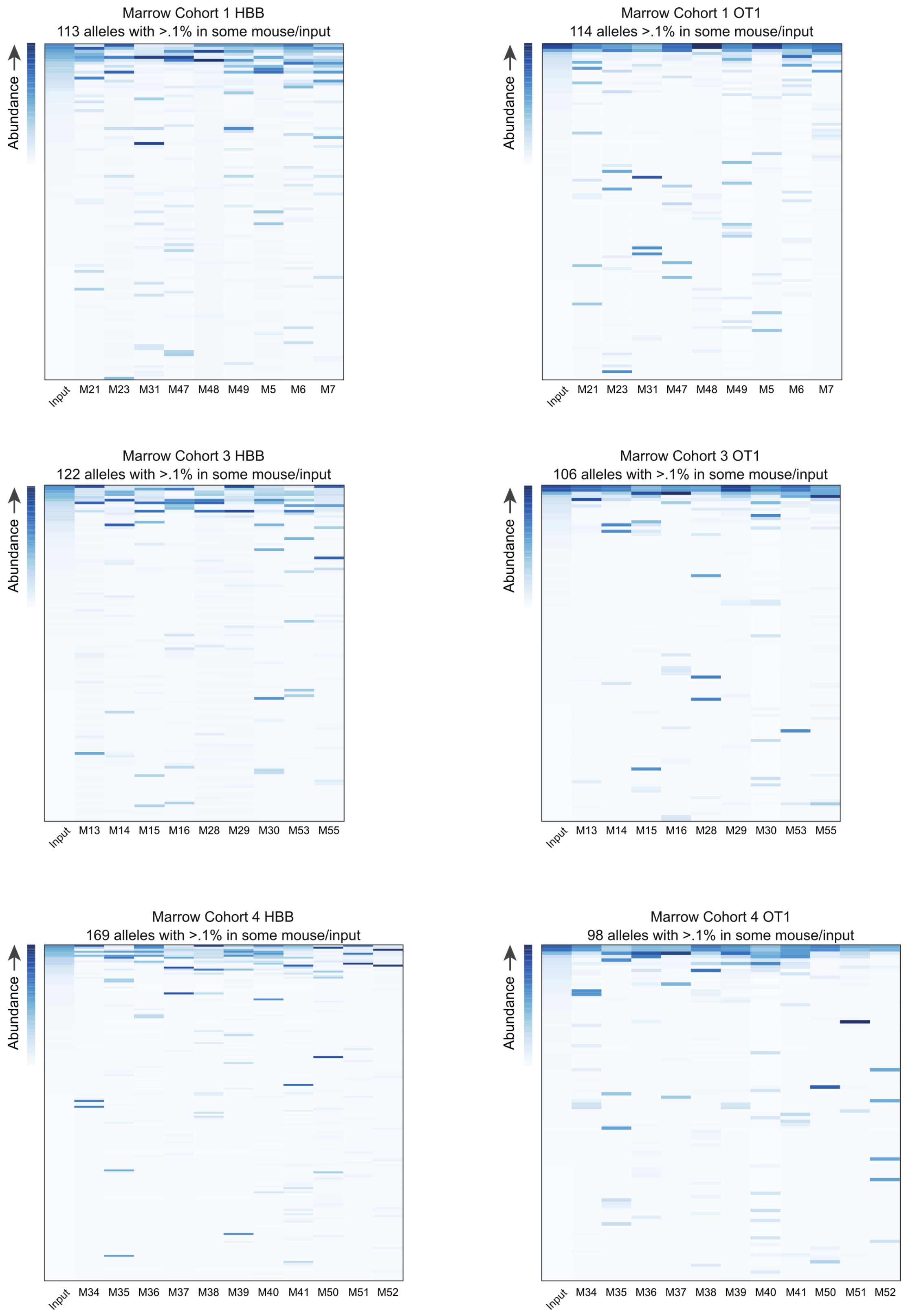

**Figure S3.** Analysis of allelic diversity in edited and xenografted hematopoietic cells: cohorts 1, 3, and 4, related to Figure 2A. Heatmaps of the 140 (*HBB*) and 135 (OT1) indel alleles whose abundance is >0.1% in at least one marrow or in the input cells. Alleles are sorted vertically by their relative frequency in the input population, and normalized in each column by frequency. Note that the most common alleles in the input population are of minor abundance in some mice, while alleles that are rare in the input may be common in one or more mice.

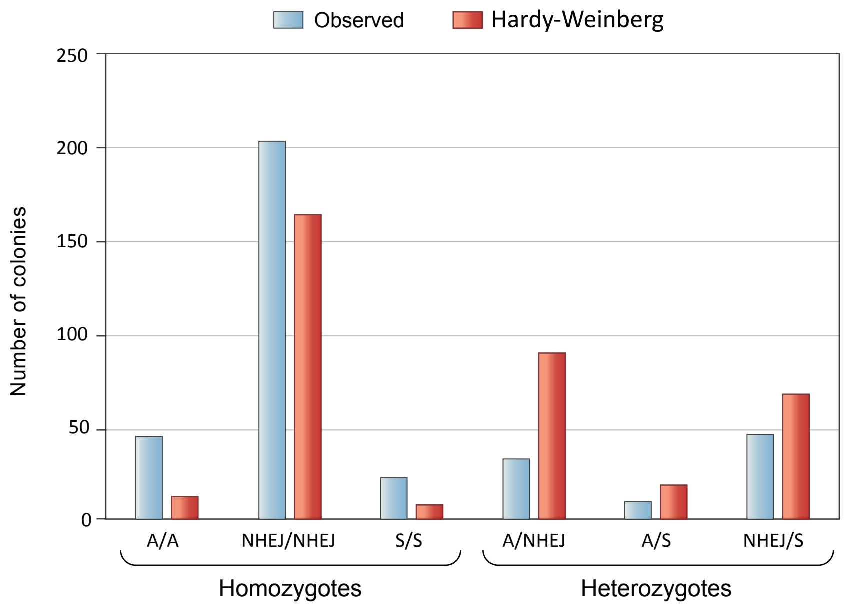

**Figure S4. *HBB* genotype frequencies in 359 clonal erythroid colonies derived from xenograft CD34+ cells isolated from mouse marrow**. The two alleles present in each clonal colony were inferred from RNA-Seq data. The y-axis shows the number of colonies with a given genotype. Alleles are identified as follows – A: homology-directed repair extends to the site of sickle mutation, producing wild-type ß-globin; NHEJ: any of 35 different alleles containing an insertion or a deletion near the CAS9 cleavage site; S: the site of sickle mutation is unaffected, comprising alleles in which homology-directed repair extended only to the PAM motif and alleles apparently unaffected by CRISPR-Cas9. Blue columns represent counts of observed genotypes, and red columns represent counts expected for allelic assortment according to the Hardy-Weinberg equilibrium (based on allele frequencies in the 359 colonies). In comparison to the distribution predicted by the Hardy-Weinberg principle, the colonies exhibit an excess of homozygous genotypes ($chi^2=10^{-37}$), with a corresponding deficit in heterozygous genotypes. This finding may reflect hemizygosity for the edited region in some apparently homozygous cells, since alleles with deletions larger than the genotyped region would not be detected by our genotyping assay. Apparent hemizygosity would result from the creation of an indel that is large enough to prevent PCR amplification and sequencing. While some colonies that are apparently homozygous for gene correction (A) may actually be heterozygous for the corrected allele and a large indel, since the wild type HBB is dominant over null (ß-thalassemia) alleles, these genotypes would not change the proportion of functionally corrected erythroid cells.

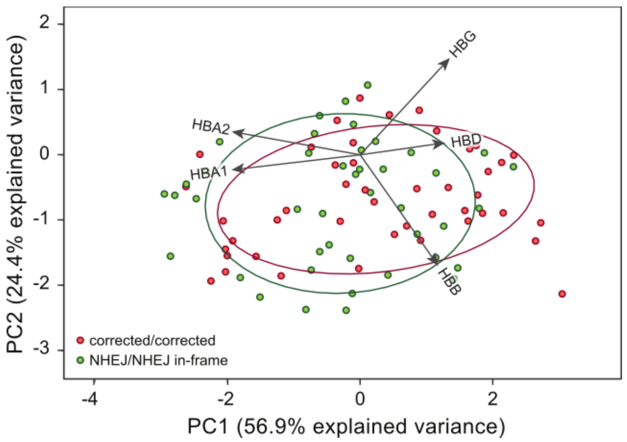

**Figure S5**. **Principal component analysis of RNA-seq data from erythroid colonies**. Comparison of colonies whose *HBB* genotypes are homozygous for in-frame indels with colonies homozygous for the corrected genotype (wild type, also shown in Figure 3C); dots represent individual colonies, and ellipses encompass 68% of colonies of each type. Colonies with in-frame indels at *HBB* have a similar ratio of *HBB* to *HBG* mRNA as corrected colonies; contrast with the out-of-frame indel colonies shown in Figure 3C.

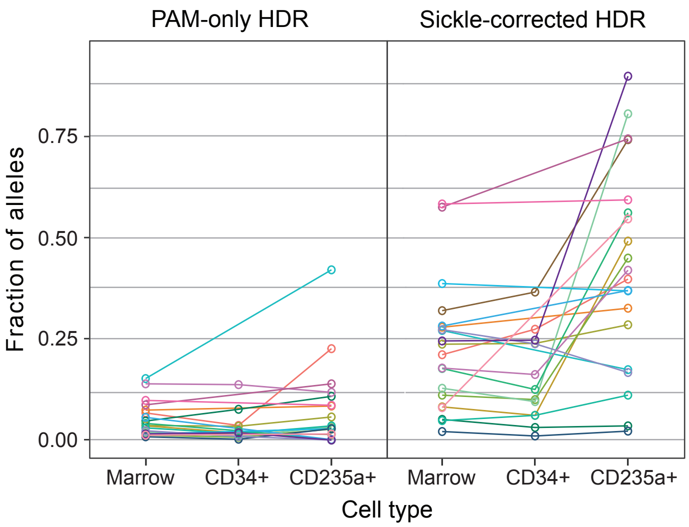

**Figure S6. Fraction of alleles that have undergone homology-directed repair (HDR) in different populations of hematopoietic cells isolated from mouse marrow: enrichment of corrected alleles in CD235+ erythroid cells**. This figure is derived from the same data as Figure 3D, but allows comparison of genotypes in different cell types from the same mouse. The left panel (PAM-only HDR) shows the fraction of haplotypes in which HDR extended to the PAM motif but not to the site of the sickle mutation; the right panel (sickle-corrected HDR) shows the fraction of haplotypes in which HDR extended to the site of the sickle mutation, correcting the sequence to wild type. Haplotype frequencies were determined in cells from whole marrow (n= 40), CD34+ cells (n=26), and CD235+ cells (n=40). Cell populations isolated from a given mouse are shown in the same color and are connected by a line.

### Wild-type Cas9 RNP (CD34+ HSPC)

| Position (1, 10, 20, 23) | Reads | |
|---|---|---|
| C C G T T A C T G C C C T G T G G G G C A A G | Reads | |
| • • • • • • • • • • • • • • • • • • • • • • • | 10776 | HBB (on-target, Chr11) |
| • • C • • • • • • • • • • • • • • • • • T G A | 2960 | OT1 intergenic Chr9 |

**Figure S7**. GUIDE-seq in CD34+ HSPC edited with the 3xMS-G10 RNP with wild-type Cas9. Only two sites, the on-target site in *HBB* and the primary off-target site OT1, are detected.

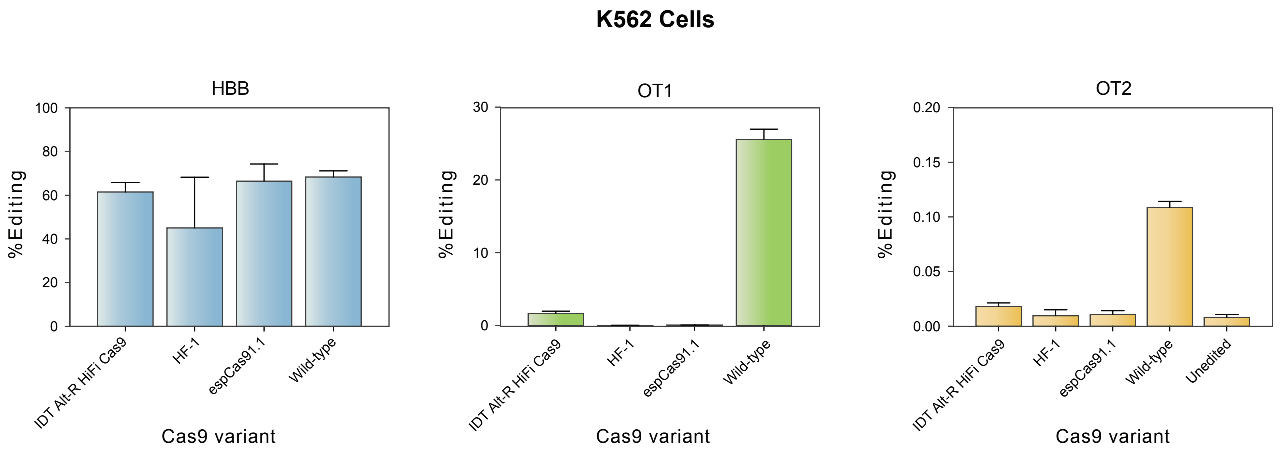

**Figure S8.** Tests of high-fidelity Cas9 RNPs in K562 cells at HBB, OT1, and OT2. Purified IDT Alt-R HiFi Cas9(*19*), Cas9 HF-1(*18*), espCas9-1.1(*17*) were complexed with guide 3xMS-G10 and delivered to K562 cells by electroporation using published methods(*10*), before analysis of editing at off-target sites OT1 and OT2 by targeted amplification (Table S3).

**Table S1**. Engraftment and editing in mice, listed by cohort and by individual mouse number.

| | | | Marrow | | CD34+ |
|---|---|---|---|---|---|
| Cohort | Mouse | % huCD45+ | % HDR | % NHEJ | % HDR |
| 1 | 5 | 89.3 | 6.256 | 79.469 | 2.528 |
| 1 | 6 | 10.8 | 29.232 | 66.908 | 21.876 |
| 1 | 7 | 9.6 | 27.016 | 68.424 | 22.108 |
| 1 | 21 | 47.2 | 19.351 | 74.06 | 15.607 |
| 1 | 23 | 60.2 | 14.576 | 80.352 | 9.428 |
| 1 | 31 | 64.1 | 28.618 | 70.545 | 34.158 |
| 1 | 47 | 70.8 | 45.959 | 32.432 | nd |
| 1 | 48 | 80.1 | 39.811 | 47.338 | nd |
| 1 | 49 | 28.2 | 26.986 | 63.094 | nd |
| 2 | 9 | 85.4 | 24.179 | 73.405 | 23.736 |
| 2 | 10 | 75.9 | 36.332 | 56.399 | 34.316 |
| 2 | 11 | 76.4 | 9.677 | 89.993 | 6.989 |
| 2 | 12 | 65.0 | 16.476 | 76.369 | 10.043 |
| 2 | 17 | 79.7 | 35.304 | 56.759 | 35.275 |
| 2 | 18 | 58.2 | 14.85 | 81.486 | 15.029 |
| 2 | 19 | 92.3 | 37.318 | 49.943 | 49.848 |
| 2 | 20 | 85.0 | 29.546 | 62.368 | 24.33 |
| 2 | 24 | 76.7 | 37.014 | 40.738 | 43.077 |
| 2 | 25 | 44.9 | 49.535 | 41.571 | 47.905 |
| 2 | 27 | 65.0 | 22.93 | 54.693 | 27.803 |
| 2 | 57 | 92.5 | 52.362 | 31.889 | nd |
| 2 | 58 | 50.7 | 51.923 | 30.47 | nd |
| 2 | 59 | 82.4 | 8.266 | 79.945 | nd |
| 3 | 13 | 25.9 | 8.987 | 89.785 | 3.476 |
| 3 | 14 | 21.2 | 47.457 | 46.969 | 55.282 |
| 3 | 15 | 63.8 | 12.403 | 86.103 | 4.556 |
| 3 | 16 | 20.7 | 8.46 | 89.226 | 3.344 |
| 3 | 28 | 76.4 | 46.094 | 46.666 | nd |
| 3 | 29 | 26.9 | 22.322 | 73.786 | 24.708 |
| 3 | 30 | 15.0 | 14.68 | 65.364 | 8.543 |
| 3 | 53 | 35.9 | 28.515 | 45.167 | 30.021 |
| 3 | 55 | 14.7 | 6.253 | 68.673 | 1.128 |
| 4 | 34 | 23.0 | 11.618 | 87.53 | 9.233 |
| 4 | 35 | 44.6 | 9.678 | 75.204 | 5.864 |
| 4 | 36 | 58.7 | 22.524 | 76.69 | 23.299 |
| 4 | 37 | 68.4 | 10.233 | 87.056 | 9.269 |
| 4 | 38 | 32.8 | 12.37 | 70.884 | 9.686 |
| 4 | 39 | 19.4 | 4.409 | 88.493 | 3.425 |
| 4 | 40 | 10.8 | 16.423 | 58.44 | 11.859 |
| 4 | 41 | 69.2 | 3.141 | 65.011 | 3.625 |
| 4 | 50 | 17.4 | 3.055 | 88.708 | 1.66 |
| 4 | 51 | 41.7 | 27.412 | 35.921 | 23.591 |
| 4 | 52 | 47.5 | 24.689 | 49.777 | 25.383 |

*nd = not determined

**Table S2**. GUIDE-seq results in K562 cells edited with either in-house produced wild-type Cas9 RNP or Alt-R HiFi Cas9 RNP, purchased from IDT, Inc. or from Aldevron Inc (sold as SpyFi Cas9).

| **Alt-R HiFi Cas9 (IDT, Inc.)** | | | | | | | | |
|---|---|---|---|---|---|---|---|---|
| Replicate 1 | | | Replicate 2 | | | Replicate 3 | | |
| Off-target site | Alias | # unique reads | Off-target site | Alias | # unique reads | | | |
| chr11:5226814 | HBB | 16891 | chr11:5226984 | HBB | 2583 | chr11:5226814 | HBB | 3789 |
| chr9:101833600 | OT1 | 425 | chr9:101833601 | OT1 | 6686 | chr9:101833600 | OT1 | 214 |
| **Alt-R HiFi Cas9 (Aldevron, Inc.)** | | | | | | | | |
| *Replicate 1* | | | *Replicate 2* | | | *Replicate 3* | | |
| Off-target site | Alias | # unique reads | Off-target site | Alias | # unique reads | Off-target site | Alias | GS reads |
| chr11:5226984 | HBB | 44802 | chr11:5226814 | HBB | 5035 | chr11:5226814 | HBB | 3522 1 |
| chr12:124319288 | OT2 | 5 | chr9:101833600 | OT1 | 665 | chr9:101833600 | OT1 | 568 |
| chr9:101833600 | OT1 | 11305 | | | | | | |
| **WT Cas9 (Berkeley)** | | | | | | | | |
| *Replicate 1* | | | *Replicate 2* | | | *Replicate 3* | | |
| Off-target site | Alias | # unique reads | Off-target site | Alias | # unique reads | Off-target site | Alias | GS reads |
| chr11:5226984 | HBB | 80313 | chr11:5226984 | HBB | 45556 | chr11:5226984 | HBB | 8998 0 |
| chr12:93155425 | | 171 | chr12:93155426 | | 88 | chr12:93155425 | | 3 |
| chr12:124319288 | OT2 | 1834 | chr12:12431929 8 | OT2 | 1718 | chr12:12431928 8 | OT2 | 1012 |
| chr17:78243561 | | 220 | chr17:68628105 | | 36 | chr12:13228822 0 | | 56 |
| chr19:33890031 | | 1532 | chr17:78243561 | | 207 | chr17:68628104 | | 56 |
| chr9:101833600 | OT1 | 49755 | chr9:101833600 | OT1 | 27211 | chr19:33890031 | | 34 |
| | | | | | | chr9:101833600 | OT1 | 5868 5 |

**Table S3**. Indel formation at GUIDE-seq, CRISTA, and CRISPor-identified off-targets analyzed using rhAmp-seq pooled primer PCR and next-generation sequencing, related to Figure 4.

| Target coordinate in hg38 | Target ID | mouse cohort 1 input | | mouse cohort 2 input | | Alt-R HiFi Cas9 HSPC | | | unedited HSPC ave | | |
|---|---|---|---|---|---|---|---|---|---|---|---|
| chr1:177624828-177624850 | | average | n | average | n | average | s.d. | n | average | s.d. | n |
| chr1:185917962-185917984 | A1_GS2 (HBB) | 90.0875 | 2 | 80.4995 | 2 | 53.13 | 2.39 | 3 | 3.34 | 3.04 | 3 |
| chr1:214841461-214841483 | A10 | 0.06 | 2 | 0.09 | 2 | 0.13 | 0.03 | 3 | 0.08 | 0.02 | 3 |
| chr1:221019077-221019099 | A11_T14 | 0.025 | 2 | 0.015 | 2 | 0.03 | 0.02 | 3 | 0.02 | 0.02 | 3 |
| chr1:223080632-223080654 | A12 | 0.125 | 2 | 0.135 | 2 | 0.09 | 0.10 | 3 | 0.14 | 0.04 | 3 |
| chr1:227859534-227859556 | A13_T150 | 0.04 | 2 | 0.045 | 2 | 0.03 | 0.01 | 3 | 0.05 | 0.01 | 3 |
| chr1:232286680-232286702 | A14 | 0.075 | 2 | 0.185 | 2 | 0.19 | 0.03 | 3 | 0.18 | 0.01 | 3 |
| chr1:242195708-242195730 | A15 | 0.035 | 2 | 0.04 | 2 | 0.02 | 0.02 | 3 | 0.03 | 0.03 | 3 |
| chr1:25195528-25195550 | A16_T148 | 0.16 | 2 | 0.115 | 2 | 0.08 | 0.04 | 3 | 0.10 | 0.07 | 3 |
| chr1:44246701-44246723 | A17_T111 | 0.075 | 2 | 0.07 | 2 | 0.07 | 0.01 | 3 | 0.07 | 0.02 | 3 |
| chr1:55821809-55821831 | A18 | 0.085 | 2 | 0.08 | 2 | 0.08 | 0.02 | 3 | 0.08 | 0.05 | 3 |
| chr1:57894707-57894729 | A19 | 0.06 | 2 | 0.095 | 2 | 0.07 | 0.01 | 3 | 0.07 | 0.02 | 3 |
| chr1:74542243-74542265 | A2_T86_GS1 (OT1) | 62.2165 | 2 | 65.657 | 2 | 2.06 | 0.40 | 3 | 0.09 | 0.01 | 3 |
| chr10:128978881-128978903 | A20_T161 | 0.085 | 2 | 0.065 | 2 | 0.03 | 0.01 | 3 | 0.03 | 0.01 | 3 |
| chr10:130898113-130898135 | A21 | 0.22 | 2 | 0.2 | 2 | 0.17 | 0.02 | 3 | 0.18 | 0.05 | 3 |
| chr10:29427735-29427757 | A22_T69 | 0.05 | 2 | 0.075 | 2 | 0.07 | 0.01 | 3 | 0.03 | 0.01 | 3 |
| chr10:57222233-57222255 | A23 | 0.045 | 2 | 0.055 | 2 | 0.06 | 0.01 | 3 | 0.07 | 0.02 | 3 |
| chr10:62728930-62728952 | A24 | 0.035 | 2 | 0.03 | 2 | 0.03 | 0.00 | 3 | 0.03 | 0.01 | 3 |
| chr10:6347856-6347878 | A25 | 0.05 | 2 | 0.02 | 2 | 0.11 | 0.02 | 2 | 0.01 | 0.02 | 3 |
| chr10:70150844-70150866 | A26_T133 | 0.16 | 2 | 0.27 | 2 | 0.25 | 0.08 | 3 | 0.17 | 0.04 | 3 |
| chr10:7041211-7041233 | A27_T71 | 0.025 | 2 | 0.03 | 2 | 0.05 | 0.03 | 3 | 0.03 | 0.02 | 3 |
| chr10:70526688-70526710 | A28_GS6 | 0.135 | 2 | 0.17 | 2 | 0.07 | 0.02 | 3 | 0.04 | 0.01 | 3 |
| chr10:71795928-71795950 | A29 | 0.04 | 2 | 0.05 | 2 | 0.03 | 0.01 | 3 | 0.07 | 0.02 | 3 |
| chr11:129972074-129972096 | A3_T146 | 0.04 | 2 | 0.035 | 2 | 0.04 | 0.02 | 3 | 0.04 | 0.02 | 3 |
| chr11:35069701-35069723 | A30_T128 | 0.35 | 2 | 0.41 | 2 | 0.43 | 0.07 | 3 | 0.31 | 0.02 | 3 |
| chr11:5226968-5226990 | A31 | 0.02 | 2 | 0.035 | 2 | 0.07 | 0.07 | 3 | 0.01 | 0.01 | 3 |
| chr11:5234379-5234402 | A32_T157 | 0.08 | 2 | 0.18 | 2 | 0.14 | 0.06 | 3 | 0.13 | 0.03 | 3 |
| chr11:5249745-5249763 | A33_T59 | 0.035 | 2 | 0.045 | 2 | 0.04 | 0.01 | 3 | 0.03 | 0.02 | 3 |
| chr11:5254687-5254705 | A34 | 0.015 | 2 | 0.025 | 2 | 0.03 | 0.02 | 3 | 0.03 | 0.02 | 3 |
| chr11:5269837-5269859 | A35_T44 | 0.79 | 2 | 1.045 | 2 | 0.48 | 0.09 | 3 | 0.70 | 0.19 | 3 |
| chr11:65628972-65628994 | A36_T6 | 0.07 | 2 | 0.055 | 2 | 0.03 | 0.01 | 3 | 0.05 | 0.05 | 3 |

| | | | | | | | | | | | |
|---|---|---|---|---|---|---|---|---|---|---|---|
| chr11:74719074-74719096 | A37_T17 | 0.06 | 2 | 0.06 | 2 | 0.07 | 0.02 | 3 | 0.05 | 0.01 | 3 |
| chr12:124319282-124319304 | A38_T31 | 0.105 | 2 | 0 | 2 | n.d. | n.d. | 0 | n.d. | n.d. | 0 |
| chr12:61898689-61898711 | A39_T105 | 0.1 | 2 | 0.145 | 2 | 0.11 | 0.04 | 3 | 0.09 | 0.04 | 3 |
| chr12:75167102-75167124 | A4_T109_GS4 (OT2) | 0.4 | 2 | 0.445 | 2 | 0.07 | 0.05 | 3 | 0.03 | 0.01 | 3 |
| chr12:93155409-93155431 | A40_T103 | 0.04 | 2 | 0.055 | 2 | 0.06 | 0.03 | 3 | 0.06 | 0.03 | 3 |
| chr13:106343233-106343255 | A41 | 0.035 | 2 | 0.045 | 2 | 0.03 | 0.02 | 3 | 0.04 | 0.02 | 3 |
| chr13:109165988-109166010 | A42_T8 | 0.09 | 2 | 0.09 | 2 | 0.10 | 0.01 | 3 | 0.07 | 0.04 | 3 |
| chr13:36881392-36881414 | A43_T79 | 0.025 | 2 | 0.03 | 2 | 0.03 | 0.01 | 3 | 0.03 | 0.01 | 3 |
| chr14:34174917-34174939 | A44 | n.d. | 0 | #DIV/0! | 0 | n.d. | n.d. | 0 | n.d. | n.d. | 0 |
| chr14:51385717-51385739 | A45 | 0.03 | 2 | 0.035 | 2 | 0.03 | 0.01 | 3 | 0.04 | 0.02 | 3 |
| chr14:72689080-72689102 | A47_T75 | 0.045 | 2 | 0.065 | 2 | 0.06 | 0.01 | 3 | 0.05 | 0.03 | 3 |
| chr14:91130157-91130179 | A48_T33 | 0.03 | 2 | 0.03 | 2 | 0.02 | 0.01 | 3 | 0.03 | 0.01 | 3 |
| chr14:94118984-94119006 | A49_T47 | 0.075 | 2 | 0.06 | 2 | 0.03 | 0.01 | 3 | 0.12 | 0.16 | 2 |
| chr14:97639186-97639208 | A5_T122 | 0.03 | 2 | 0.06 | 2 | 0.05 | 0.01 | 3 | 0.05 | 0.01 | 3 |
| chr15:46305914-46305936 | A50_T50 | 0.05 | 2 | 0.08 | 2 | 0.04 | 0.03 | 3 | 0.02 | 0.01 | 3 |
| chr15:78827238-78827260 | A51 | 0.115 | 2 | 0.11 | 2 | 0.15 | 0.10 | 3 | 0.12 | 0.06 | 3 |
| chr15:80912338-80912360 | A52_T100 | 0.105 | 2 | 0.16 | 2 | 0.11 | 0.03 | 3 | 0.13 | 0.06 | 3 |
| chr15:87945584-87945606 | A53_T93 | 0.16 | 2 | 0.095 | 2 | 0.12 | 0.01 | 3 | 0.11 | 0.03 | 3 |
| chr15:99995162-99995184 | A54 | 0.035 | 2 | 0.06 | 2 | 0.07 | 0.02 | 3 | 0.07 | 0.02 | 3 |
| chr16:19723993-19724015 | A6_T140_GS5 | 0.115 | 2 | 0.095 | 2 | 0.05 | 0.02 | 3 | 0.06 | 0.03 | 3 |
| chr16:29744860-29744882 | A7_T108 | 0.18 | 2 | 0.075 | 2 | 0.06 | 0.02 | 3 | 0.03 | 0.01 | 3 |
| chr16:48451185-48451207 | A8_T23 | 0.02 | 2 | 0.025 | 2 | 0.03 | 0.01 | 3 | 0.03 | 0.00 | 3 |
| chr16:56253545-56253567 | A9_T89 | 0.115 | 2 | 0.13 | 2 | 0.11 | 0.04 | 3 | 0.11 | 0.02 | 3 |
| chr16:67969032-67969055 | aav1 | 0.045 | 2 | 0.07 | 2 | 0.07 | 0.03 | 3 | 0.06 | 0.01 | 3 |
| chr16:73389721-73389743 | GS3 | 0.135 | 2 | 0.145 | 2 | 0.16 | 0.02 | 3 | 0.09 | 0.08 | 3 |
| chr16:85326988-85327010 | ssDNA1 | 0.16 | 2 | 0.165 | 2 | 0.15 | 0.02 | 3 | 0.16 | 0.03 | 3 |
| chr16:8663652-8663674 | ssDNA2 | 0.015 | 2 | 0.02 | 2 | 0.02 | 0.01 | 3 | 0.02 | 0.01 | 3 |
| chr16:90037480-90037502 | ssDNA4 | 0.205 | 2 | 0.155 | 2 | 0.14 | 0.02 | 3 | 0.20 | 0.14 | 3 |
| chr17:16858685-16858707 | ssDNA5 | 0.235 | 2 | 0.395 | 2 | 0.44 | 0.09 | 3 | 0.38 | 0.16 | 3 |
| chr17:17017415-17017437 | T1 | 0.05 | 2 | 0.075 | 2 | 0.07 | 0.03 | 3 | 0.09 | 0.03 | 3 |
| chr17:17445739-17445761 | T10 | 0.53 | 2 | 0.535 | 2 | 0.55 | 0.02 | 3 | 0.50 | 0.07 | 3 |
| chr17:18401409-18401431 | T101 | 0.2 | 2 | 0.18 | 2 | 0.16 | 0.02 | 3 | 0.18 | 0.04 | 3 |
| chr17:20543009-20543031 | T102 | 0.01 | 2 | 0.01 | 2 | 0.01 | 0.01 | 3 | 0.01 | 0.01 | 3 |
| chr17:20569256-20569278 | T104 | 0.03 | 2 | 0.03 | 2 | 0.03 | 0.02 | 3 | 0.04 | 0.02 | 3 |
| chr17:33734299-33734321 | T106 | n.d. | 0 | #DIV/0! | 0 | n.d. | n.d. | 0 | n.d. | n.d. | 0 |
| chr17:36211903-36211925 | T107 | 0.265 | 2 | 0.73 | 2 | 0.60 | 0.37 | 3 | 0.49 | 0.18 | 3 |

| | | | | | | | | | | | |
|---|---|---|---|---|---|---|---|---|---|---|---|
| chr17:40491755-40491778 | T11 | 0.095 | 2 | 0.07 | 2 | 0.09 | 0.02 | 3 | 0.08 | 0.02 | 3 |
| chr17:44078405-44078427 | T110 | 0.135 | 2 | 0.14 | 2 | 0.05 | 0.01 | 3 | 0.06 | 0.04 | 3 |
| chr17:5138723-5138745 | T112 | 0.125 | 2 | 0.1 | 2 | 0.10 | 0.01 | 3 | 0.08 | 0.02 | 3 |
| chr17:68628098-68628120 | T113 | 0.05 | 2 | 0.06 | 2 | 0.12 | 0.03 | 3 | 0.12 | 0.02 | 3 |
| chr17:73257465-73257487 | T114 | 0.1 | 2 | 0.07 | 2 | 0.07 | 0.01 | 3 | 0.08 | 0.02 | 3 |
| chr17:78243543-78243567 | T115 | 0.22 | 2 | 0.17 | 2 | 0.21 | 0.02 | 3 | 0.17 | 0.03 | 3 |
| chr18:10140361-10140383 | T116 | 0.025 | 2 | 0.055 | 2 | 0.05 | 0.00 | 3 | 0.06 | 0.04 | 3 |
| chr18:4126020-4126042 | T117 | 0.04 | 2 | 0.035 | 2 | 0.03 | 0.01 | 3 | 0.05 | 0.02 | 3 |
| chr18:45734262-45734284 | T118 | 0.095 | 2 | 0.105 | 2 | 0.09 | 0.03 | 3 | 0.08 | 0.03 | 3 |
| chr18:6663828-6663850 | T119 | 0.025 | 2 | 0.035 | 2 | 0.03 | 0.01 | 3 | 0.04 | 0.02 | 3 |
| chr18:77451781-77451803 | T12 | 0.095 | 2 | 0.08 | 2 | 0.07 | 0.02 | 3 | 0.08 | 0.02 | 3 |
| chr19:33890015-33890037 | T120 | 0.095 | 2 | 0.085 | 2 | 0.07 | 0.01 | 3 | 0.06 | 0.01 | 3 |
| chr19:35738742-35738764 | T121 | 0.06 | 2 | 0.14 | 2 | 0.10 | 0.02 | 3 | 0.08 | 0.03 | 3 |
| chr19:37048134-37048156 | T123 | 0.055 | 2 | 0.06 | 2 | 0.05 | 0.01 | 3 | 0.06 | 0.01 | 3 |
| chr19:48630752-48630774 | T124 | 0.075 | 2 | 0.085 | 2 | 0.10 | 0.11 | 3 | 0.12 | 0.11 | 3 |
| chr19:48638948-48638970 | T125 | 0.015 | 2 | 0.02 | 2 | 0.02 | 0.02 | 3 | 0.02 | 0.01 | 3 |
| chr19:49506736-49506758 | T126 | 0.03 | 2 | 0.04 | 2 | 0.03 | 0.01 | 3 | 0.05 | 0.02 | 3 |
| chr2:111258773-111258795 | T127 | 0.05 | 2 | 0.07 | 2 | 0.05 | 0.03 | 3 | 0.08 | 0.04 | 3 |
| chr2:11637669-11637691 | T129 | 0.05 | 2 | 0.075 | 2 | 0.09 | 0.06 | 3 | 0.09 | 0.08 | 3 |
| chr2:119659823-119659845 | T13 | 0.02 | 2 | 0.025 | 2 | 0.01 | 0.01 | 3 | 0.04 | 0.02 | 3 |
| chr2:120957647-120957669 | T130 | 0.04 | 2 | 0.04 | 2 | 0.04 | 0.01 | 3 | 0.03 | 0.02 | 3 |
| chr2:122461575-122461597 | T131 | 0.01 | 2 | 0.035 | 2 | 0.02 | 0.02 | 3 | 0.02 | 0.02 | 3 |
| chr2:162706706-162706728 | T132 | 0.07 | 2 | 0.095 | 2 | 0.06 | 0.01 | 3 | 0.05 | 0.01 | 3 |
| chr2:171602229-171602251 | T134 | 0.035 | 2 | 0.05 | 2 | n.d. | n.d. | 0 | n.d. | n.d. | 0 |
| chr2:211341537-211341559 | T135 | 0.05 | 2 | 0.045 | 2 | 0.05 | 0.01 | 3 | 0.05 | 0.03 | 3 |
| chr2:218065329-218065351 | T136 | 0.055 | 2 | 0.04 | 2 | 0.07 | 0.02 | 3 | 0.04 | 0.03 | 3 |
| chr2:224133104-224133126 | T137 | 0.245 | 2 | 0.24 | 2 | 0.25 | 0.04 | 3 | 0.24 | 0.02 | 3 |
| chr2:229415994-229416016 | T138 | n.d. | 0 | #DIV/0! | 0 | n.d. | n.d. | 0 | n.d. | n.d. | 0 |
| chr2:237377872-237377894 | T139 | 0.045 | 2 | 0.07 | 2 | 0.04 | 0.02 | 3 | 0.04 | 0.02 | 3 |
| chr2:239053899-239053921 | T141 | 0.28 | 2 | 0.29 | 2 | 0.30 | 0.02 | 3 | 0.30 | 0.02 | 3 |
| chr2:239712022-239712044 | T142 | 0.115 | 2 | 0.115 | 2 | 0.13 | 0.03 | 3 | 0.10 | 0.02 | 3 |
| chr2:240968117-240968139 | T143 | 0.13 | 2 | 0.135 | 2 | 0.09 | 0.02 | 3 | 0.13 | 0.03 | 3 |
| chr2:29883206-29883228 | T144 | 0.045 | 2 | 0.045 | 2 | 0.03 | 0.01 | 3 | 0.04 | 0.01 | 3 |
| chr2:43273462-43273484 | T145 | 0.08 | 2 | 0.09 | 2 | 0.07 | 0.01 | 3 | 0.07 | 0.02 | 3 |
| chr2:79966568-79966590 | T147 | 0.29 | 2 | 0.255 | 2 | 0.26 | 0.05 | 3 | 0.22 | 0.04 | 3 |
| chr2:87692118-87692140 | T149 | 0.05 | 2 | 0.085 | 2 | n.d. | n.d. | 0 | n.d. | n.d. | 0 |
| chr20:35227470-35227493 | T15 | 0.065 | 2 | 0.08 | 2 | 0.07 | 0.03 | 3 | 0.06 | 0.03 | 3 |
| chr20:59561148-59561170 | T151 | 0.215 | 2 | 0.205 | 2 | 0.18 | 0.01 | 3 | 0.16 | 0.04 | 3 |
| chr22:16006635-16006657 | T152 | 0.08 | 2 | 0.075 | 2 | 0.06 | 0.03 | 3 | 0.08 | 0.03 | 3 |

| | | | | | | | | | | | |
|---|---|---|---|---|---|---|---|---|---|---|---|
| chr22:16045816-16045838 | T153 | n.d. | 0 | #DIV/0! | 0 | n.d. | n.d. | 0 | n.d. | n.d. | 0 |
| chr22:16749716-16749738 | T154 | 0.05 | 2 | 0.155 | 2 | 0.12 | 0.03 | 3 | 0.09 | 0.03 | 3 |
| chr22:35141396-35141418 | T155 | 0.115 | 2 | 0.1 | 2 | 0.09 | 0.03 | 3 | 0.12 | 0.02 | 3 |
| chr22:37182645-37182667 | T156 | 0.115 | 2 | 0.11 | 2 | 0.11 | 0.04 | 3 | 0.11 | 0.04 | 3 |
| chr3:10396267-10396289 | T158 | 0.11 | 2 | 0.155 | 2 | 0.14 | 0.02 | 3 | 0.14 | 0.04 | 3 |
| chr3:111251318-111251340 | T159 | 0.06 | 2 | 0.075 | 2 | 0.06 | 0.03 | 3 | 0.06 | 0.02 | 3 |
| chr3:119034494-119034516 | T16 | 0.105 | 2 | 0.145 | 2 | 0.12 | 0.02 | 3 | 0.09 | 0.01 | 3 |
| chr3:128394820-128394842 | T160 | 0.03 | 2 | 0.045 | 2 | 0.03 | 0.03 | 3 | 0.03 | 0.03 | 3 |
| chr3:13877218-13877240 | T162 | 0.14 | 2 | 0.1 | 2 | 0.08 | 0.02 | 3 | 0.08 | 0.03 | 3 |
| chr3:14448080-14448102 | T18 | 0.02 | 2 | 0.02 | 2 | 0.03 | 0.02 | 3 | 0.01 | 0.01 | 3 |
| chr3:150727088-150727110 | T19 | 0.025 | 2 | 0.035 | 2 | 0.03 | 0.01 | 3 | 0.03 | 0.00 | 3 |
| chr3:156019243-156019265 | T2 | 0.05 | 2 | 0.07 | 2 | 0.09 | 0.02 | 3 | 0.12 | 0.02 | 3 |
| chr3:167309719-167309741 | T20 | 0.07 | 2 | 0.055 | 2 | 0.06 | 0.02 | 3 | 0.08 | 0.02 | 3 |
| chr3:181316329-181316351 | T21 | 0.06 | 2 | 0.08 | 2 | 0.06 | 0.01 | 3 | 0.08 | 0.02 | 3 |
| chr3:40750370-40750392 | T22 | 0.04 | 2 | 0.045 | 2 | 0.05 | 0.02 | 3 | 0.03 | 0.02 | 3 |
| chr3:50315247-50315269 | T24 | 0.05 | 2 | 0.05 | 2 | 0.06 | 0.01 | 3 | 0.06 | 0.04 | 3 |
| chr3:56928987-56929009 | T25 | 0.105 | 2 | 0.09 | 2 | 0.14 | 0.09 | 3 | 0.11 | 0.03 | 3 |
| chr4:1004154-1004176 | T26 | 0.15 | 2 | 0.055 | 2 | 0.03 | 0.01 | 3 | 0.04 | 0.03 | 3 |
| chr4:107044371-107044393 | T27 | 0.07 | 2 | 0.075 | 2 | 0.10 | 0.03 | 3 | 0.08 | 0.04 | 3 |
| chr4:1777993-1778015 | T28 | 0.045 | 2 | 0.06 | 2 | 0.06 | 0.03 | 3 | 0.07 | 0.04 | 3 |
| chr4:54236432-54236454 | T29 | 0.11 | 2 | 0.105 | 2 | 0.08 | 0.01 | 3 | 0.09 | 0.01 | 3 |
| chr4:54254203-54254225 | T3 | 0.155 | 2 | 0.225 | 2 | 0.12 | 0.03 | 3 | 0.10 | 0.04 | 3 |
| chr4:94367421-94367443 | T30 | n.d. | 0 | #DIV/0! | 0 | n.d. | n.d. | 0 | n.d. | n.d. | 0 |
| chr5:102752174-102752196 | T32 | 0.11 | 2 | 0.105 | 2 | 0.11 | 0.03 | 3 | 0.10 | 0.02 | 3 |
| chr5:129539830-129539852 | T34 | 0.03 | 2 | 0.03 | 2 | 0.03 | 0.01 | 3 | 0.03 | 0.02 | 3 |
| chr5:132087686-132087708 | T35 | 0.02 | 2 | 0.02 | 2 | 0.03 | 0.02 | 3 | 0.04 | 0.02 | 3 |
| chr5:134545147-134545169 | T36 | 0.055 | 2 | 0.055 | 2 | 0.04 | 0.02 | 3 | 0.04 | 0.03 | 3 |
| chr5:14346925-14346947 | T37 | 0.085 | 2 | 0.075 | 2 | 0.04 | 0.03 | 3 | 0.06 | 0.03 | 3 |
| chr5:163951815-163951837 | T38 | 1.19 | 2 | 1.37 | 2 | 0.67 | 0.09 | 3 | 0.77 | 0.02 | 3 |
| chr5:167881725-167881747 | T39 | 0.05 | 2 | 0.045 | 2 | 0.04 | 0.02 | 3 | 0.04 | 0.01 | 3 |
| chr5:171688565-171688587 | T4 | 0.05 | 2 | 0.035 | 2 | 0.04 | 0.01 | 3 | 0.05 | 0.03 | 3 |
| chr5:176728638-176728660 | T40 | 0.04 | 2 | 0.055 | 2 | 0.05 | 0.02 | 3 | 0.07 | 0.01 | 3 |
| chr5:180133430-180133452 | T41 | 0.115 | 2 | 0.125 | 2 | 0.10 | 0.02 | 3 | 0.11 | 0.03 | 3 |
| chr5:44465642-44465664 | T42 | 0.11 | 2 | 0.11 | 2 | 0.11 | 0.02 | 3 | 0.13 | 0.00 | 3 |
| chr6:149230734-149230756 | T43 | 0.235 | 2 | 0.29 | 2 | 0.22 | 0.15 | 3 | 0.31 | 0.19 | 3 |
| chr6:156050368-156050390 | T45 | 0.13 | 2 | 0.17 | 2 | 0.16 | 0.02 | 3 | 0.18 | 0.03 | 3 |
| chr6:156836330-156836352 | T46 | 0.06 | 2 | 0.065 | 2 | 0.06 | 0.02 | 3 | 0.06 | 0.01 | 3 |
| chr6:3088439-3088461 | T48 | 0.105 | 2 | 0.08 | 2 | 0.07 | 0.00 | 3 | 0.06 | 0.01 | 3 |
| chr6:31724781-31724803 | T49 | 0.16 | 2 | 0.15 | 2 | 0.12 | 0.02 | 3 | 0.11 | 0.03 | 3 |

| | | | | | | | | | | | |
|---|---|---|---|---|---|---|---|---|---|---|---|
| chr6:34561693-34561715 | T5 | 0.055 | 2 | 0.05 | 2 | 0.04 | 0.01 | 3 | 0.05 | 0.03 | 3 |
| chr6:42245244-42245266 | T51 | 0.17 | 2 | 0.18 | 2 | 0.24 | 0.04 | 3 | 0.22 | 0.02 | 3 |
| chr6:43287221-43287243 | T52 | 0.085 | 2 | 0.075 | 2 | 0.08 | 0.02 | 3 | 0.09 | 0.04 | 3 |
| chr6:50073642-50073664 | T53 | 0.27 | 2 | 0.175 | 2 | 0.20 | 0.02 | 3 | 0.23 | 0.04 | 3 |
| chr6:71331681-71331703 | T54 | 0.115 | 2 | 0.125 | 2 | 0.12 | 0.01 | 3 | 0.09 | 0.03 | 3 |
| chr6:82973754-82973776 | T55 | 0.03 | 2 | 0.045 | 2 | 0.03 | 0.01 | 3 | 0.03 | 0.02 | 3 |
| chr6:95813282-95813304 | T56 | 0.09 | 2 | 0.08 | 2 | 0.08 | 0.02 | 3 | 0.09 | 0.04 | 3 |
| chr7:104843096-104843118 | T57 | 0.035 | 2 | 0.045 | 2 | 0.05 | 0.03 | 3 | 0.04 | 0.03 | 3 |
| chr7:114806995-114807017 | T58 | 0.055 | 2 | 0.035 | 2 | 0.03 | 0.03 | 3 | 0.03 | 0.02 | 3 |
| chr7:127598434-127598456 | T60 | 0.1 | 2 | 0.07 | 2 | 0.06 | 0.02 | 3 | 0.07 | 0.03 | 3 |
| chr7:151451690-151451712 | T61 | 0.04 | 2 | 0.05 | 2 | 0.05 | 0.02 | 3 | 0.04 | 0.02 | 3 |
| chr7:155442737-155442759 | T62 | 0.07 | 2 | 0.08 | 2 | 0.08 | 0.02 | 3 | 0.08 | 0.01 | 3 |
| chr7:158362273-158362295 | T63 | 0.04 | 2 | 0.03 | 2 | 0.03 | 0.01 | 3 | 0.04 | 0.01 | 3 |
| chr7:1901770-1901801 | T64 | 0.08 | 2 | 0.085 | 2 | 0.09 | 0.06 | 3 | 0.09 | 0.04 | 3 |
| chr7:45586859-45586881 | T65 | 0.09 | 2 | 0.105 | 2 | 0.09 | 0.01 | 3 | 0.13 | 0.07 | 3 |
| chr7:74538053-74538075 | T66 | 0.07 | 2 | 0.065 | 2 | 0.07 | 0.04 | 3 | 0.06 | 0.01 | 3 |
| chr7:76263475-76263497 | T67 | 0.11 | 2 | 0.05 | 2 | 0.09 | 0.02 | 3 | 0.08 | 0.02 | 3 |
| chr7:7854915-7854937 | T68 | 0.07 | 2 | 0.06 | 2 | 0.11 | 0.04 | 3 | 0.08 | 0.02 | 3 |
| chr7:94349301-94349323 | T7 | 0.1 | 2 | 0.11 | 2 | 0.11 | 0.02 | 3 | 0.11 | 0.03 | 3 |
| chr7:98244842-98244864 | T70 | 0.085 | 2 | 0.115 | 2 | 0.13 | 0.02 | 3 | 0.10 | 0.03 | 3 |
| chr8:101473699-101473721 | T72 | 0.03 | 2 | 0.04 | 2 | 0.03 | 0.02 | 3 | 0.03 | 0.01 | 3 |
| chr8:133012196-133012218 | T73 | 0.03 | 2 | 0.045 | 2 | 0.03 | 0.00 | 3 | 0.05 | 0.02 | 3 |
| chr8:133349316-133349338 | T74 | 0.065 | 2 | 0.075 | 2 | 0.08 | 0.04 | 3 | 0.08 | 0.01 | 3 |
| chr8:143801773-143801795 | T76 | 0.195 | 2 | 0.19 | 2 | 0.18 | 0.02 | 3 | 0.16 | 0.03 | 3 |
| chr8:21771827-21771849 | T77 | 0.07 | 2 | 0.06 | 2 | 0.04 | 0.02 | 3 | 0.06 | 0.08 | 3 |
| chr8:22098771-22098793 | T78 | 0.055 | 2 | 0.09 | 2 | 0.06 | 0.01 | 3 | 0.07 | 0.02 | 3 |
| chr8:39934669-39934691 | T80 | 0.085 | 2 | 0.13 | 2 | 0.14 | 0.02 | 3 | 0.10 | 0.04 | 3 |
| chr9:101833584-101833606 | T81 | 0.02 | 2 | 0.04 | 2 | 0.03 | 0.01 | 3 | 0.03 | 0.01 | 3 |
| chr9:105198188-105198210 | T82 | 0.385 | 2 | 0.315 | 2 | 0.31 | 0.02 | 3 | 0.32 | 0.02 | 3 |
| chr9:11915772-11915794 | T83 | 0.04 | 2 | 0.06 | 2 | 0.03 | 0.02 | 3 | 0.05 | 0.02 | 3 |
| chr9:131734280-131734302 | T84 | 0.155 | 2 | 0.175 | 2 | 0.16 | 0.02 | 3 | 0.13 | 0.02 | 3 |
| chr9:132119560-132119582 | T85 | 0.16 | 2 | 0.13 | 2 | n.d. | n.d. | 0 | n.d. | n.d. | 0 |
| chr9:132637599-132637621 | T87 | 0.29 | 2 | 0.315 | 2 | 0.34 | 0.03 | 3 | 0.30 | 0.05 | 3 |
| chr9:136562377-136562399 | T88 | 0.17 | 2 | 0.2 | 2 | 0.18 | 0.07 | 3 | 0.17 | 0.03 | 3 |
| chr9:137546904-137546926 | T9 | 0.1 | 2 | 0.135 | 2 | 0.14 | 0.05 | 3 | 0.11 | 0.09 | 3 |
| chr9:34198705-34198727 | T90 | 0.045 | 2 | 0.05 | 2 | 0.06 | 0.02 | 3 | 0.05 | 0.02 | 3 |
| chr9:42191521-42191543 | T91 | 0.115 | 2 | 0.125 | 2 | 0.12 | 0.03 | 3 | 0.11 | 0.03 | 3 |
| chr9:4421255-4421277 | T92 | 0.09 | 2 | 0.06 | 2 | 0.06 | 0.01 | 3 | 0.10 | 0.06 | 3 |
| chrX:100296997-100297019 | T94 | 0.05 | 2 | 0.065 | 2 | 0.07 | 0.03 | 3 | 0.06 | 0.01 | 3 |

| | | | | | | | | | | | |
|---|---|---|---|---|---|---|---|---|---|---|---|
| chrX:38926721-38926743 | T95 | 0.125 | 2 | 0.16 | 2 | 0.14 | 0.01 | 3 | 0.11 | 0.04 | 3 |
| chrX:4589543-4589565 | T96 | 0.235 | 2 | 0.225 | 2 | 0.26 | 0.02 | 3 | 0.12 | 0.01 | 3 |
| chrX:68998829-68998851 | T97 | 0.035 | 2 | 0.03 | 2 | 0.05 | 0.02 | 3 | 0.04 | 0.03 | 3 |
| chrX:75786405-75786427 | T98 | 0.055 | 2 | 0.025 | 2 | 0.03 | 0.03 | 3 | 0.02 | 0.01 | 3 |
| | T99 | 0.08 | 2 | 0.08 | 2 | 0.11 | 0.05 | 3 | 0.12 | 0.11 | 3 |
